# A lactate-dependent shift of glycolysis mediates synaptic and cognitive processes

**DOI:** 10.1101/2023.03.15.532748

**Authors:** Ignacio Fernández-Moncada, Gianluca Lavanco, Unai B. Fundazuri, Nasrin Bollmohr, Sarah Mountadem, Pauline Hachaguer, Francisca Julio-Kalajzic, Doriane Gisquet, Tommaso Dalla Tor, Roman Serrat, Luigi Bellocchio, Astrid Cannich, Bérénice Fortunato-Marsol, Yusuke Nasu, Robert E. Campbell, Filippo Drago, Carla Cannizzaro, Guillaume Ferreira, Anne-Karine Bouzier-Sore, Luc Pellerin, Juan P. Bolaños, Gilles Bonvento, L. Felipe Barros, Stephane H. R. Oliet, Aude Panatier, Giovanni Marsicano

**Author notes:** These authors contributed equally (Gianluca Lavanco, Unai B. Fundazuri, Nasrin Bollmohr, Sarah Mountadem). These authors share senior authorship (Aude Panatier, Giovanni Marsicano). Correspondence (G.M.), (I.F-M.).

## Abstract

Control of brain energy metabolism and regulation of synaptic activity through gliotransmission are two important ways, through which astrocytes contribute to mental functions. However, the potential functional and molecular links between these two astrocyte-dependent processes have been scantly explored. Here we show that a lactate-dependent shift of glycolysis underlies the production of the gliotransmitter D-serine by acute activation of astrocyte type-1 cannabinoid (CB1) receptors, thereby gating synaptic and cognitive processes. Acute cannabinoid application causes a CB1 receptor-dependent rapid and reversible increase of lactate production and release in primary astrocyte cultures. As shown before, mutant mice lacking the CB1 receptor gene in astrocytes (GFAP-CB1-KO) were impaired in a novel object recognition (NOR) memory task. This phenotype was rescued not only by the gliotransmitter D-serine, but also by its precursor L-serine. Surprisingly, the administration of lactate and of an agonist of the lactate receptor HCAR1 also reverted the memory impairment of GFAP-CB1-KO mice. This rescue effect was abolished by *in vivo* blockade of the astrocyte-specific phosphorylated pathway (PP), which diverts glycolysis towards L-serine synthesis, suggesting that lactate signaling might promote the accumulation of this amino acid. Consistent with this idea, lactate and HCAR1 agonism increased the co-agonist occupancy of CA1 post-synaptic hippocampal NMDA receptors. This effect of lactate was abolished by blockade of PP. By establishing a mechanistic link between lactate production and signaling, serine availability, synaptic activity and behavior, these results reveal an unforeseen functional connection between energy metabolism and gliotransmission to control cognitive processes.

## MAIN

Type-1 cannabinoid (CB1) receptors are G-protein coupled receptors (GPCRs) prominently expressed across the central nervous system^1–3^. Physiological engagement of CB1 receptors by their endogenous ligands, the endocannabinoids, modulates many behavioral processes, including food intake, emotional responses and cognition^4^. Importantly, CB1 receptors can also be targeted by exogenous cannabinoids, such as Δ^9^-tetrahydrocannabinol (THC), the main psychoactive component of *Cannabis sativa*. This exogenous, non-physiological activation might bear therapeutic properties, but it can also alter brain activity, impairing, for instance, cognitive, locomotor and perceptive functions^5^. Another remarkable feature of CB1 receptor signaling is its subcellular compartmentalization, which deviates from the strict plasma membrane functional localization of most GPCRs. Thus, few but functionally significant brain CB1 receptors are found in association with mitochondrial membranes (mtCB1 receptors), where they can alter mitochondrial functions and regulate behavior^6–10^. This uncommon subcellular distribution allows CB1 receptors modulating brain functions *via* parallel signaling pathways triggered by specific subcellular pools^11^.

CB1 receptors are highly expressed in neurons, but they are also present at low but functionally very relevant levels in other brain cell types, such as astrocytes^12^. Notably, astroglial CB1 receptors can govern brain functions and cognitive processes, such as novel object recognition (NOR)^13, 14^. Recently, it has been shown that astrocytes also possess functional mtCB1 receptors^8, 15, 16^. In particular, persistent (24 hours) activation of mtCB1 receptors in astrocytes results in decreased mitochondrial functions and diminished lactate production. This astroglial metabolic failure brings about neuronal stress and impairment of social behavior as observed 24 hours after cannabinoid exposure^16^.

The present study originated from the idea of detailing the specific molecular underpinnings of such negative mtCB1 receptor-dependent control of brain lactate levels. However, early experiments lead to the surprising observation that short-term exposure to cannabinoid agonists can rapidly, reliably and transiently increase lactate levels in astrocytes. Therefore, we set off to investigate the molecular mechanisms and the relevance of this observation. The results revealed an unexpected molecular link between metabolic and signaling properties of astrocytes, which can regulate physiological cognitive processes.

## RESULTS

As previously shown using other methods^16^, 24-hour stimulation of CB1 receptors in cultured astrocytes expressing the reporter Laconic (Ref. 17) led to a reduction of intracellular lactate levels (Extended Data Fig. 1A,B). With the intention to investigate the temporal progression of this effect, we then analyzed the short-term impact of cannabinoid treatment. To our surprise, we observed that acute application of the CB1 agonist WIN55,212-2 (WIN55) was able to rapidly *increase* lactate levels in cultured astrocytes (Extended Data Fig. 1C,D). Therefore, we decided to investigate the molecular mechanisms and the potential behavioral relevance of this phenomenon from the physiological and pharmacological points of view.

Parallel cultures of astrocytes from wild-type (CB1-WT) and mutant CB1-KO mice expressing the reporter Laconic were shortly exposed to WIN55 (1 µM) and the fluorescent responses were imaged and quantified. We observed that the transient intracellular lactate increase induced by WIN55 was fully dependent on the presence of the CB1 receptor (Fig. 1A-C). The previously described negative effect of persistent CB1 agonism on lactate levels depends on mtCB1 receptors^16^. Thus, we tested if this specific subcellular pool was also involved in the short-term lactate increase induced by WIN55. Remarkably, this effect did not require mtCB1 receptors, as the lactate rise was not altered in astrocytes derived from DN22-CB1-KI mice, a knock-in line in which the DN22-CB1 protein (a mutant version of CB1 lacking mitochondrial localization^7^) replaces the wild-type protein^7, 18^ (Fig. 1A-C). Of note, the basal levels of lactate or capacity to produce lactate were not altered by the genotype of the astrocytes (Extended data Fig. 2A-C). Moreover, blockade of mitochondrial oxidative phosphorylation (OXPHOS) with sodium azide triggered lactate increases with similar amplitude and kinetics in CB1-WT, CB1-KO and DN22-CB1-KI astrocytes (Extended data Fig. 2D-I). These data indicates that the differential acute effects of WIN55 on these cells could not be ascribed to differences in their basal levels of lactate or to their general ability to produce or accumulate this metabolite.

**Figure 1.**
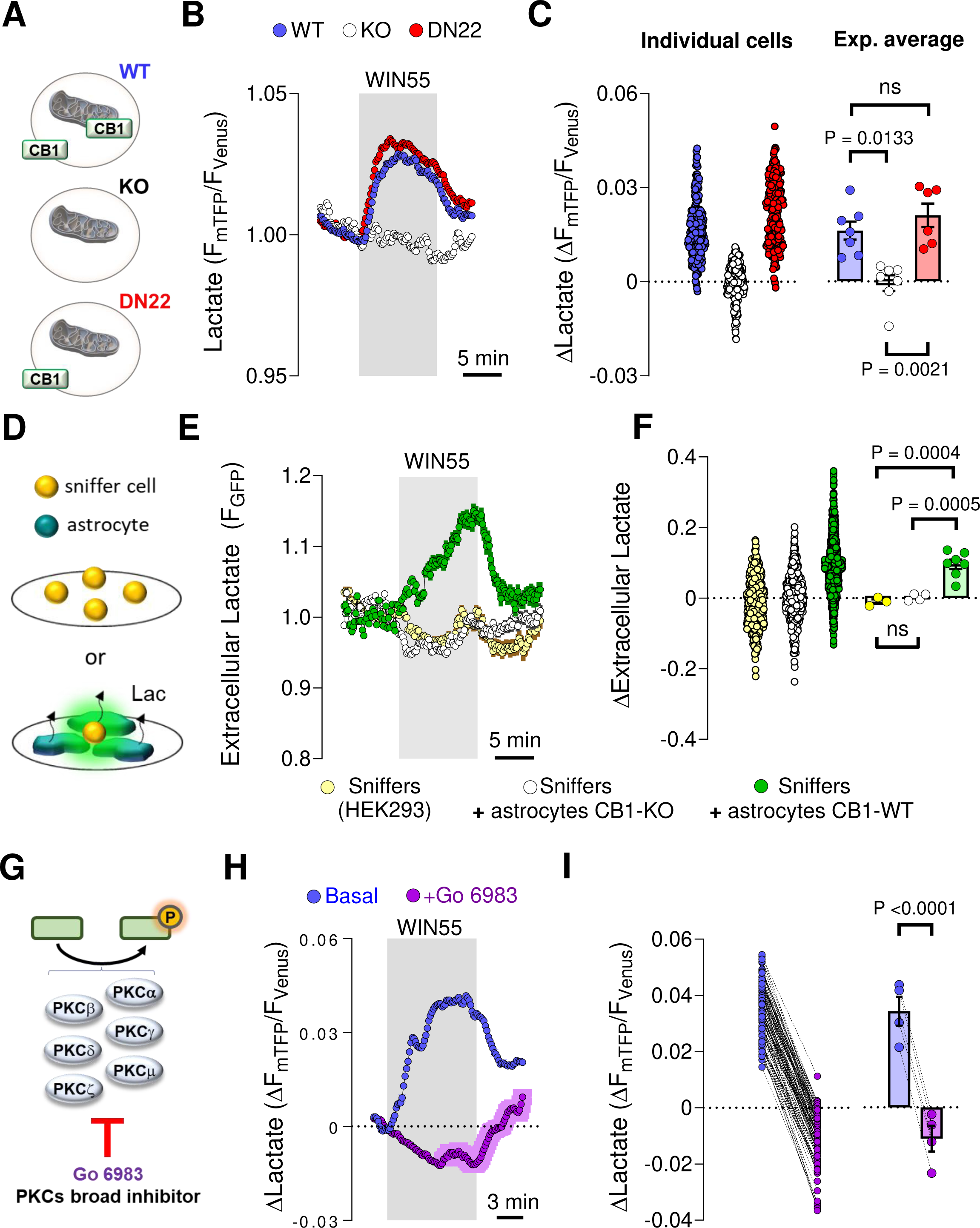
Acute stimulation of astrocytes lactate metabolism by CB1 receptors. **A**, Subcellular localization of CB1 receptors in the primary astrocytes culture models. **B**, Intracellular lactate measurement during exposure to WIN55 (1 μM), in CB1-WT (blue circles, average of 42 cells), CB1-KO (white circles, average of 29 cells) and DN22-CB1-KI (red circles, average of 42 cells) astrocytes. **C**, Quantification of lactate changes after 5 min exposure to WIN55. CB1-WT: n = 7, 247 cells. CB1-KO: n = 7, 252 cells. DN22-CB1-KI: n=6, 235 cells. **D**, Sniffer cell strategy for determination of extracellular lactate levels. HEK cells expressing an extracellular lactate fluorescent biosensor (sniffers cells) were cultured alone or in co-culture with astrocytes. **E**, Extracellular lactate measurements during exposure to WIN55 (1 μM), in a pure culture of sniffers cells (yellow circles, average of 81 cells), a co-culture of sniffers cells and CB1-WT astrocytes (green circles, average of 76 cells) and a co-culture of sniffers cells and CB1-KO astrocytes (white circles, average of 102 cells). **F**, Quantification of the extracellular lactate levels after 10 min exposure to WIN55. Sniffer culture: n=3, 406 cells. Sniffer + CB1-WT astrocytes co-culture: n = 7, 997 cells. Sniffer + CB1-KO astrocytes co-culture: n = 4, 469 cells. **G**, Scheme depicting the PKC isozymes inhibited by Go 6983. **H**, Intracellular lactate measurement during exposure to WIN55 (1 μM), in CB1-WT astrocytes, before (blue) and during exposure to the broad PKC inhibitor Go 6983 (purple). N = 4, 124 cells. **I**, Quantification of lactate changes after 5 min exposure to WIN55, before (blue) and during exposure to Go 6983 (purple), n = 4, 124 cells analyzed. Data corresponds to the average (mean±SEM) of representative of a single (**B**,**E**) or all experiments (**G**). Circles in scatter or before-after plots correspond to individual cells (**C**,**F**,**I**). Bars correspond to experiments average (mean±SEM), with circles representing individual experiment average (**C**,**F**,**I**). Statistical analysis was performed using a Kruskal-Wallis test followed by Dunn’s multiple comparison test (**A**), One-way ANOVA followed by Tukey’s multiple comparison test (**F**) and two-tailed paired t-test (**I**). See Supplementary Table 2 for more details.

Intracellular increases of lactate can be due to enhanced production, but also to decreased release. To dissect these components in the acute effects of cannabinoid on lactate dynamics, we measured astrocyte lactate production *via* a transport-stop technique^17, 19^. The effect of the broad monocarboxylate transporter (MCT) inhibitor Diclofenac ^20, 21^ on lactate accumulation is fully reversible (Extended data Fig. 3A,B). This allows devising a paired assessment of lactate production by measuring the rate of accumulation upon MCT block, before and during WIN55 application (Extended data Fig. 3C). Stimulation of CB1 receptors in WT astrocytes resulted in a significant increase in the rate of intracellular lactate accumulation, indicating augmented lactate production (Extended data Fig. 3C,D). To exert its physiological functions, lactate is generally extruded from astrocytes into the extracellular space^22, 23^. Thus, we next asked whether the WIN55-induced lactate production was accompanied by increased release of the metabolite. To explore this possibility, we adapted a “sniffer cells” strategy^24, 25^, in which HEK cells expressing an extracellular lactate fluorescent biosensor^26^ are able to detect the amount of ambient lactate levels in an extracellular medium (Fig. 1D), in the presence of a constant buffer superfusion. Whereas WIN55 did not alter the extracellular lactate levels in a pure culture of sniffer cells, its application to a co-culture of sniffer cells with CB1-WT astrocytes led to an extracellular lactate accumulation (Fig. 1E,F). Importantly, extracellular lactate remained unchanged upon WIN55 exposure when sniffers cells were mixed with CB1-KO astrocytes (Fig. 1E,F).

Next, we asked what molecular mechanisms might participate in the astrocyte CB1 receptor-mediated lactate increase. The protein kinase C (PKC) has been shown to be transiently activated by CB1 receptor activation and to mediate short-term amnesic effects of cannabinoids^27^. After verifying that the magnitudes of sequential WIN55-induced lactate increases are similar (Extended data Fig. 4A-B), we quantified the effect of WIN55 on lactate levels before and during exposure to Go 6983, a broad pharmacological blocker of PKC activity^28, 29^ (Fig. 1G,H and Extended data Fig. 4C). Interestingly, the inhibitor completely abolished the lactate increase induced by WIN55 (Fig. 1H,I and Extended data Fig. 4C).

Thus, opposite to the persistent negative effects involving mitochondrial CB1 receptor signaling^16^, short-term activation of non-mitochondrial associated astroglial CB1 receptors results into the PKC-dependent transient stimulation of lactate production and release.

To determine if the quick stimulation of lactate metabolism mediated by astroglial CB1 receptors is relevant for brain functions, we took advantage of the known role of endocannabinoid signaling in the long-term memory version of the novel object recognition (NOR) task^14^. Mice lacking CB1 receptors in cells expressing the astrocyte marker glial fibrillary acidic protein (GFAP-CB1-KO mice)^30^ are impaired in long-term NOR performance^14^. This phenotype has been explained by impairment of hippocampal synaptic D-serine availability and consequent impairment of synaptic N-Methyl-D-Aspartate Receptors (NMDAR) functions during the consolidation phase of this task^14^. Noteworthy, it is not known if this physiological control of NOR performance depends on the mitochondrial pool of astroglial CB1 receptors. To address this point, we used a specific double-viral rescue approach to delete astroglial CB1 receptors and re-express either the CB1-WT or the DN22-CB1 proteins in the hippocampus of CB1-floxed mice^8, 11, 14, 31^, thereby generating Control, HPC-GFAP-CB1-KO, HPC-GFAP-CB1-RS and HPC-GFAP-DN22-CB1-RS mice, respectively (see Methods, Supplementary Table 1 and Fig. 2A). As expected^14^, the deletion of CB1 receptors from hippocampal astrocytes resulted in impaired NOR performance (Fig. 2B). Remarkably, this impairment of HPC-GFAP-CB1-KO mice was rescued by re-expression of both wild-type CB1 and mutant DN22-CB1 in HPC-GFAP-CB1-RS and HPC-GFAP-DN22-CB1-RS, respectively (Fig. 2B and Extended data Fig. 5). This indicates that mtCB1 receptor signaling is not necessary for physiological endocannabinoid-dependent control of NOR performance.

**Figure 2.**
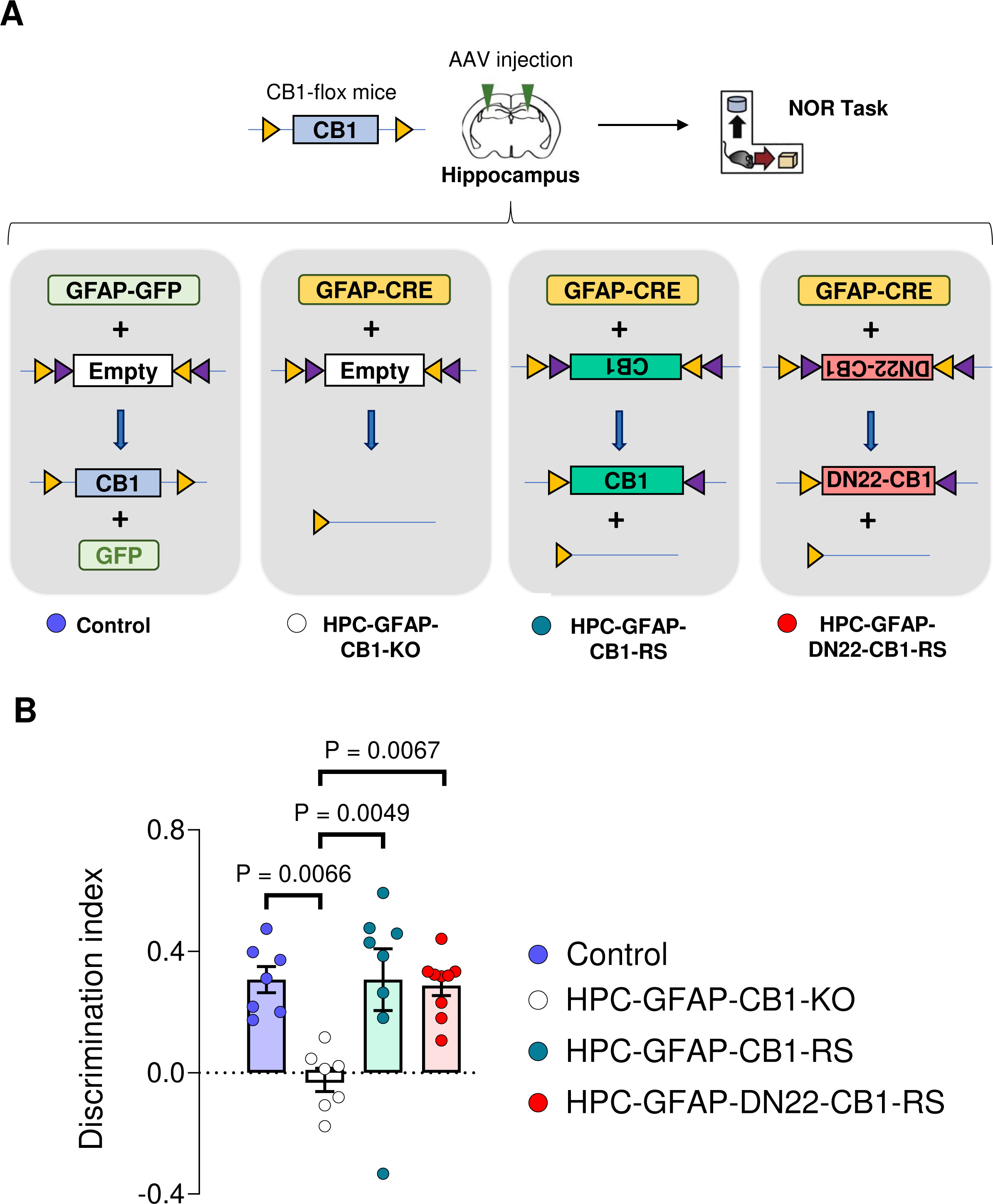
The astroglial mtCB1 receptor is not necessary for physiological NOR performance. **A**, Schematic of the viral approach used for deletion of hippocampal astroglial CB1 receptors and expression rescue with either wild type or DN22-CB1 sequences. Four groups of animals; Control, HPC-GFAP-CB1-KO, HPC-GFAP-CB1-RS and HPC-GFAP-DN22-CB1-RS mice; were obtained by *s*tereotactic injection of a mix of specific AAV constructs (see material and methods for details). **B**, Summary of NOR performance in Control (blue), HPC-GFAP-CB1-KO (white), HPC-GFAP-CB1-RS (teal) and HPC-GFAP-DN22-CB1-RS (red) mice, n= 7-9 mice per condition. Bars correspond to experiments average (mean±SEM) and circles represent individual animals (**B**). Statistical analysis was performed using a One-way ANOVA followed by Tukey’s multiple comparison test (**B**). See Supplementary Table 2 for more details.

The data collected so far show that non-mitochondrial astroglial CB1 receptors can both increase lactate accumulation and mediate physiological NOR performance. Considering that efficient lactate metabolism is required for several behavioral processes^22, 32^, we asked whether this metabolic function of astroglial CB1 receptors might contribute to determining the synaptic activity required for NOR memory consolidation. A post-training intraperitoneal (I.P.) injection of lactate at a concentration known to reach the brain parenchyma (1 g/kg)^33^, was able to fully rescue the NOR impairment of GFAP-CB1-KO mice (Fig. 3A and Extended data Fig. 6A). Next, we asked whether this rescuing effect was due to the energetic or signaling properties of lactate in the brain^22, 34^. To disentangle this point, we tested if the hydroxycarboxylic acid receptor 1 (HCAR1)^35, 36^ is involved in this effect of lactate. An IP injection of the HCAR1 agonist 3,5-dihydroxybenzoic acid (3,5-DHBA; 240 mg/kg)^37^ had no effect on wild-type animals, but it fully rescued the NOR impairment of GFAP-CB1-KO littermates (Fig. 3A and Extended data Fig. 6A), indicating that the effect of lactate is likely due to activation of its cognate receptor. As both lactate and 3,5-DHBA rescuing effects were very similar to the one obtained with D-serine^14^, we hypothesized that astroglial CB1 receptor control of lactate signaling might participate in the regulation of synaptic D-serine levels to provide physiological NOR performance. D-serine is an amino acid derived from L-serine^38–41^, which, in the adult brain, is mainly produced by astrocytes *via* the consumption of the glycolytic intermediate 3-phosphoglycerate (3PG) in the phosphorylated pathway^38^ (Fig. 3B). To test whether astroglial CB1 receptor-dependent increase of lactate might impact L-serine activity in the brain, we first assessed the potential impact of L-serine on astroglial CB1 receptor-dependent NOR performance. An I.P. injection of L-serine (0.5 g/kg) was also able to rescue the memory deficit of GFAP-CB1-KO mice (Fig. 3A and Extended data Fig. 6A), suggesting that the impaired D-serine availability in these mutants^14^ might be ascribed to a decreased astroglial L-serine production. To address this idea and identify the potential relationship between lactate and serine signaling, we adopted a pharmacological approach to inhibit phosphoglycerate dehydrogenase (PHGDH), the enzyme providing the first step of L-serine production in the phosphorylated pathway^38^ (Fig. 3B). The administration of high doses of the PHGDH blocker NCT-503 (Fig. 3C)^42^ alone impaired long-term NOR memory consolidation, possibly due to direct inhibition of L- and D-serine availability (Extended data Fig. 6B,C). Thus, we performed a full dose-response study to identify a sub-effective dose of NCT-503 that does not alter long-term memory formation *per se* (Extended data Fig. 6B,C). Then, we tested the ability of lactate or 3,5-DHBA to rescue the memory impairment of GFAP-CB1-KO mice under vehicle or in the presence of 6 mg/kg NCT-503. This treatment did not alter the NOR performance of GFAP-CB1-WT mice (P=0.8037, compare Fig. 3A and 3D; Supplementary Table 2) or the impaired memory of their GFAP-CB1-KO littermates (Fig. 3D and Extended data Fig. 7D). Conversely, the drug fully abolished both the lactate- and 3,5-DHBA-induced rescue of NOR performance of GFAP-CB1-KO mice (Fig. 3D and Extended data Fig. 7D). Importantly, the delivery of L-serine was still able to rescue the long-term memory impairment in the presence of the same dose of NCT-503 (Fig. 3D and Extended data Fig. 7D), indicating that the action of L-serine is downstream of the PHGDH activity.

**Figure 3.**
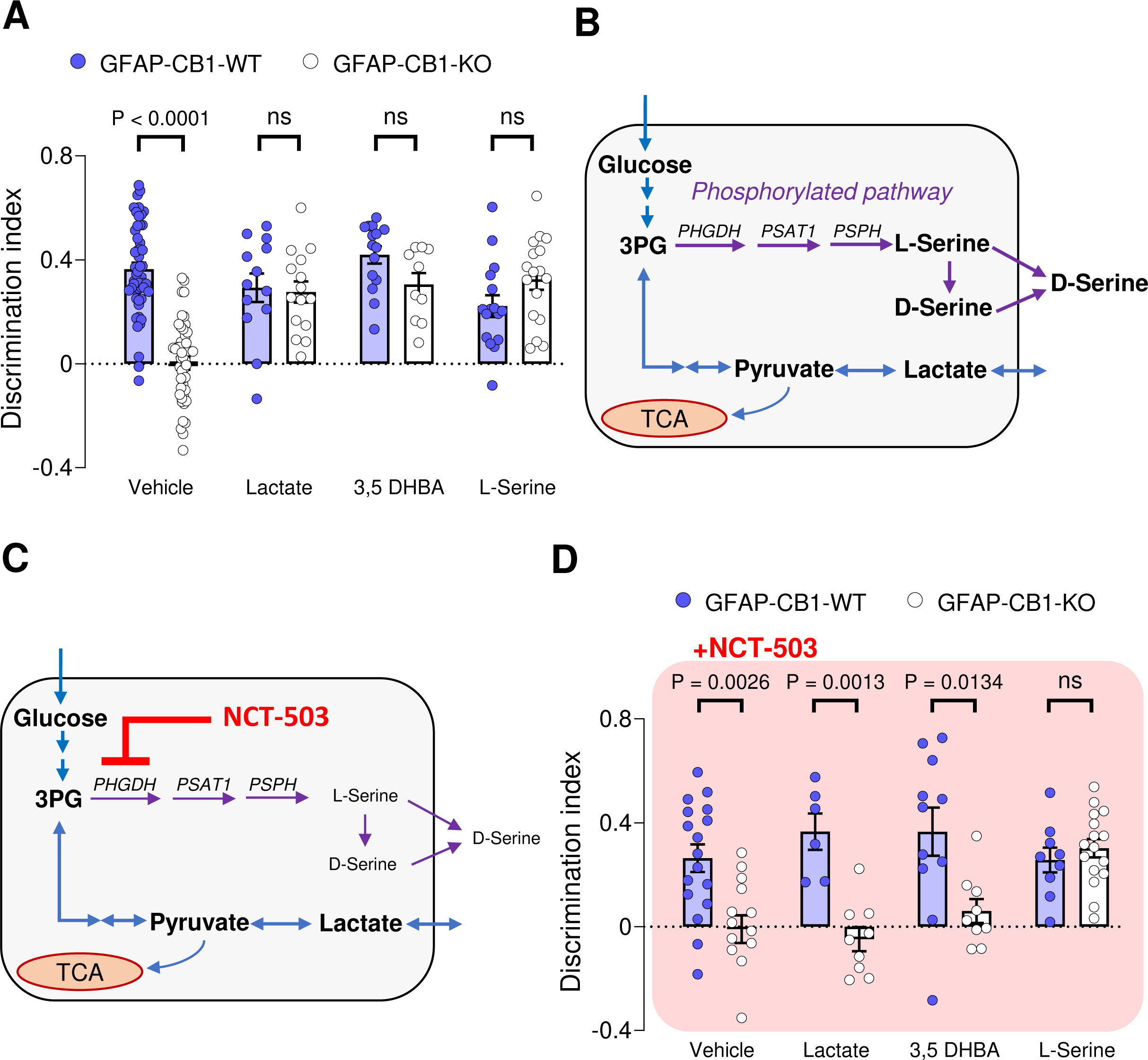
Lactate promotes NOR performance *via* receptor-dependent stimulation of the phosphorylated pathway. **A**, NOR performance of GFAP-CB1-WT (blue circles) and GFAP-CB1-KO (white circles) mice. Animals were treated either with an I.P. injection of vehicle, lactate (1 g/kg), 3,5-DHBA (240 mg/kg) or L-serine (0.5 g/kg), immediately after the acquisition phase. N= 18-53 mice per condition. **B**, Scheme depicting the lactate production and the phosphorylated pathway, and its interaction at the level of 3-phosphoglycerate (3PG). **C**, Scheme depicting the expected effect of a sub-effective dose of NCT-503 on the L-serine synthesis. **D**, NOR performance of GFAP-CB1-WT (blue circles) and GFAP-CB1-KO (white circles) mice. Animals were treated either with an I.P. injection of vehicle + NCT-503 (6 mg/kg), lactate (1 g/kg) + NCT-503, 3,5-DHBA (240 mg/kg) + NCT-503 or L-serine (0.5 g/kg) + NCT-503, immediately after the acquisition phase. N= 6-20 mice per condition. Bars correspond to experiments average (mean±SEM) and circles represent individual animals (**A**,**D**). Statistical analysis was performed using a two-way ANOVA followed by a Tukey’s multiple comparison test (**A**,**D**). See Supplementary Table 2 for more details.

Overall, these results suggest that the physiological activation of astroglial CB1 receptors enables cognitive performance *via* stimulation of glucose metabolism, increase of lactate supply, activation of HCAR1 signaling and potentiation of L- and D-serine availability *via* the phosphorylated pathway.

These results suggest that lactate might control synaptic D-serine availability, eventually resulting in the adequate signaling of NMDARs. However, it is still possible that the mechanisms underlying the rescue effect of lactate in the NOR performance of GFAP-CB1-KO mice are explained by compensatory mechanisms developed under the specific conditions of the mutant mice. In other words, the deletion of the CB1 gene in astrocytes might induce alterations that provide lactate with functions that it does not have under physiological conditions. To address this point and to clarify the role of lactate levels on the dynamics of synaptic D-serine availability, we explored if the link between these metabolites as observed in the GFAP-CB1-KO mice is also present in WT animals. To analyze the synaptic D-serine availability, we performed electrophysiological extracellular field recordings (fEPSPs) of synaptic NMDARs in the *stratum radiatum* of CA1 area in wild-type hippocampal slices^14, 41, 43^ (NMDAR-fEPSPs, Extended data Fig. 7A). In control conditions, the co-agonist binding site occupancy of synaptic NMDARs are not fully saturated, thus exogenous bath application of D-serine (50 µM) results in the increase of synaptic NMDAR activity and therefore an increase of NMDAR-fEPSPs slope^14, 41, 43^ (Fig. 4A). Intriguingly, bath application of exogenous lactate (2 mM) was able to potentiate NMDAR activity with a magnitude similar to the one induced by exogenous D-serine (Fig. 4A). However, this lactate-induced potentiation was slower than the one triggered by D-serine (Fig. 4B). To test whether this lactate effect was downstream the increase in D-serine availability, the co-agonist binding sites of synaptic NMDARs were first saturated with exogenous D-serine (50 µM) and then lactate was bath applied. Notably, lactate application had no impact on the slope of NMDAR-fEPSPs and therefore on synaptic NMDAR activity in these conditions (Fig. 4C), suggesting that the potentiating effect of the metabolite might be due to a downstream increase in D-serine availability. However, it is still possible that the application of D-serine might cause a “ceiling effect”, impeding a serine-independent effect of lactate to be observed. Therefore, we directly tested whether the activity of the phosphorylated pathway was necessary for the potentiation on synaptic NMDAR activity by lactate. Strikingly, this effect was blunted in slices preincubated with NCT-503 (Fig. 4D and Extended data Fig. 7B), showing that PHGDH activity is a required step of the process. Of note, the potentiation induced by D-serine was not altered by NCT-503 (Extended data Fig. 7C), further suggesting that lactate plays an upstream role in the astrocyte cascade leading to increased D-serine availability at synaptic NMDARs. Finally, we tested whether HCAR1 signaling was involved in the lactate-mediated control of synaptic D-serine availability. Bath application of 3,5-DHBA (1 mM) resulted in the increase of synaptic NMDARs activity (Fig. 4E). However, and importantly, prior application of 3,5-DHBA (1 mM) resulted in a complete occlusion of the exogenous D-serine effect on NMDAR activity (Fig. 4F), indicating that the activation of HCAR1 promotes an increase in synaptic D-serine availability. These results confirm that lactate can modulate the phosphorylated pathway *via* activation of HCAR1 signaling to control synaptic D-serine availability independently of CB1 receptor genetic deletion, supporting a physiological role for this phenomenon.

**Figure 4.**
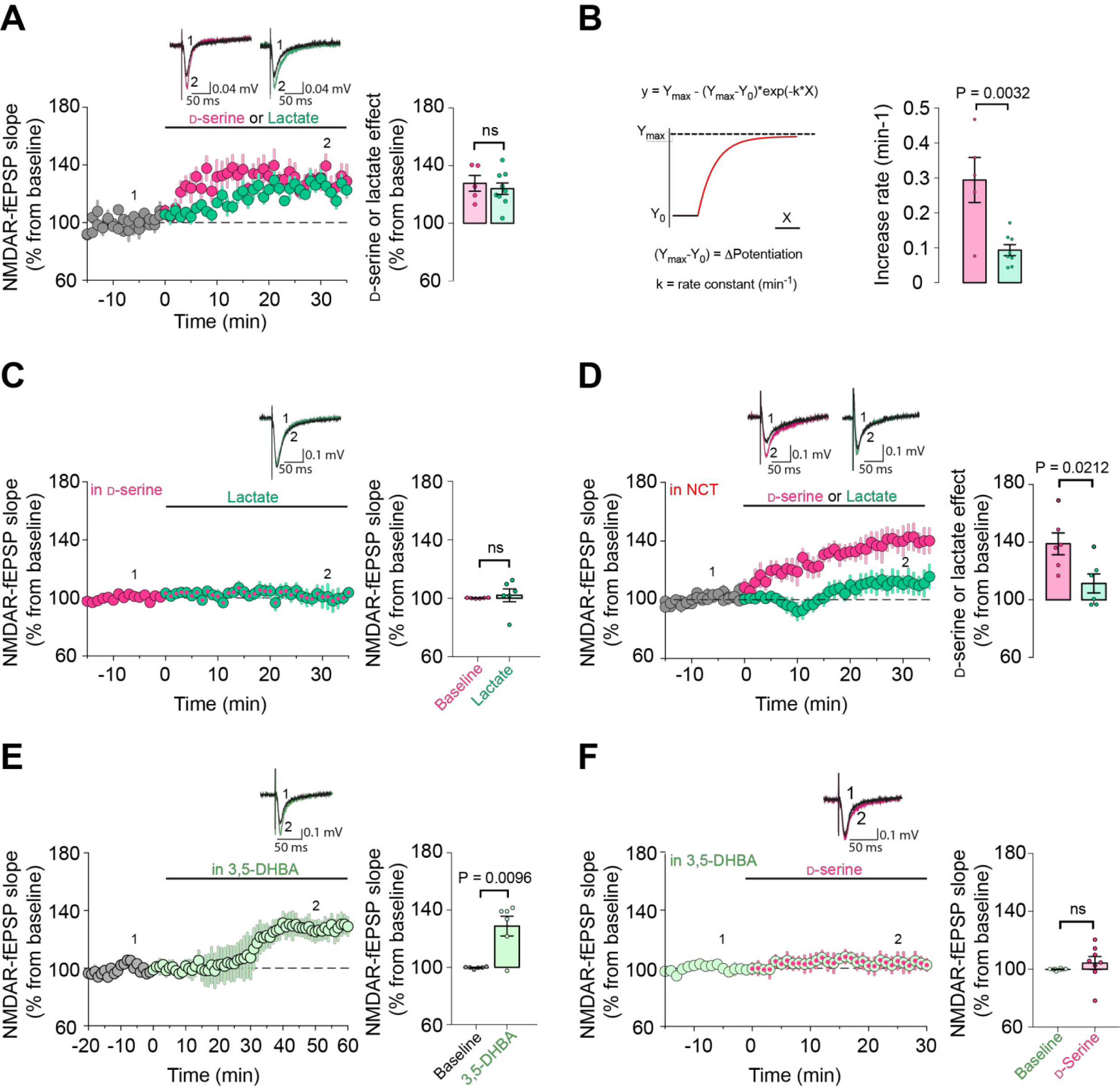
Synaptic D-serine availability is modulated by lactate. **A**, On the top, representative averaged NMDAR-fEPSPs traces from 30 consecutives sweeps evoked (1) 10 min before and (2) during (25 to 35 min) bath application of D-serine (magenta) or lactate (green). The summary plots show the effect of exogenous D-serine (50 μM – magenta circles, n=5 slices, 4 mice) or lactate (2 mM – green circles, n=9 slices, 8 mice) on NMDAR-mediated fEPSPs slopes in acute WT hippocampal slices. **B**, Non-linear model and equation used (left) to fit the data shown in panel A, to determine the increase rate (right) of the NMDAR-mediated fEPSPs potentiation induced by lactate or D-serine. **C,** On the top, representative averaged NMDAR-fEPSPs traces evoked (1) 10 min before and (2) during (25 to 35 min) bath application of lactate (green) in the presence of D-serine. The summary plots show the NMDAR-fEPSPs slopes before (magenta) and during lactate exposure (2 mM – magenta/green, n=6 slices, 6 mice) in slices preincubated with D-serine (50 μM). **D**, On the top, representative averaged NMDAR-fEPSPs traces evoked (1) 10 min before and (2) during (25 to 35 min) bath application of D-serine (in magenta) or lactate (in green) in the presence of NCT-503. The summary plots show the effect of exogenous D-serine (50 μM – magenta, n=6 slices, 6 mice) or lactate (2 mM – green, n=6 slices, 6 mice) on NMDAR-mediated fEPSPs slopes in slices preincubated with NCT-503 (10-20 μM). **E**, On the top, representatives NMDAR-fEPSPs traces evoked (1) 10 min before and (2) during bath application (25 to 35 min) of 3,5-DHBA (light green). Summary plots showing the effect of 3,5-DHBA (1 mM– light green circles, n=6 slices, 4 mice) on NMDAR-mediated fEPSPs slopes. **F**, On the top, representatives NMDAR-fEPSPs traces evoked (1) before and (2) during bath application of exogenous D-serine in the presence of 3,5-DHBA. The summary plots show the NMDAR-fEPSPs slopes before (light green) and during exogenous D-serine exposure (50 μM – light green/magenta circles, n=8 slices, 5 mice) in slices preincubated with 3,5-DHBA (1 mM). Data are presented as mean±SEM (**A**,**C-F**). Bars correspond to experiments average (mean±SEM) and circles represent single measurement (**A**-**F**). Statistical analysis was performed using a two-tailed unpaired t-test (**A**-**F**). See Supplementary Table 2 for more details.

The data described so far are compatible with a scenario, in which acute activation of hippocampal astroglial CB1 receptors leads to increased lactate production and release. This lactate would in turn promote L- and D-serine signaling at NMDARs *via* activation of HCAR1, eventually mediating consolidation of NOR memory. This physiological scenario seems at odds with the previously described pharmacological effects of CB1 receptor activation^7, 16, 44^. Indeed, *in vivo* activation of CB1 receptor by drugs does not only reduce lactate levels^16^, but it also impairs long-term NOR memory^7, 44^. Therefore, we wondered what could be the mechanistic explanation of these opposite effects of endogenous and pharmacological activation of CB1 receptors. The major differences between endogenous and exogenous CB1 receptor agonists rely on their spatiotemporal features. Specifically, whereas physiological production of endocannabinoids is generally spatially restricted and it is followed by rapid degradation^1, 45, 46^, exogenous cannabinoids are likely to unselectively spread throughout the body and their action is temporally limited only by their pharmacokinetic properties^1^. In other words, the effects of physiologically produced endocannabinoids are generally very local and short lasting, whereas the effects of exogenously administered CB1 receptor agonists are global and long lasting. Based on this reasoning, we asked what is the time course and the mechanisms possibly mediating the switch between the CB1 receptor-dependent increase (present data) and decrease of lactate (Ref 16) in cultured astrocytes. A 70 minutes-long application of WIN55 to astrocyte cultures expressing the Laconic lactate sensor resulted in a temporal biphasic effect, with the acute increase of lactate occurring during the first 5-10 minutes and a later shift towards a decrease, which became significant at 60-70 minutes after application (Fig. 5A and Extended data Fig. 8A). We previously showed that the decrease of lactate observed after 24-hours cannabinoid incubation requires mtCB1 receptors^16^. Consistently, 70 minutes treatment of astrocytes derived from DN22-CB1-KI mice with WIN55 resulted in an increase of lactate, which never shifted to a decrease (Fig. 5B and Extended data Fig. 8B). Altogether, these data indicate that the respective mtCB1 receptor-independent increase and mtCB1 receptor-dependent decrease of lactate levels in astrocytes are a function of time.

**Figure 5.**
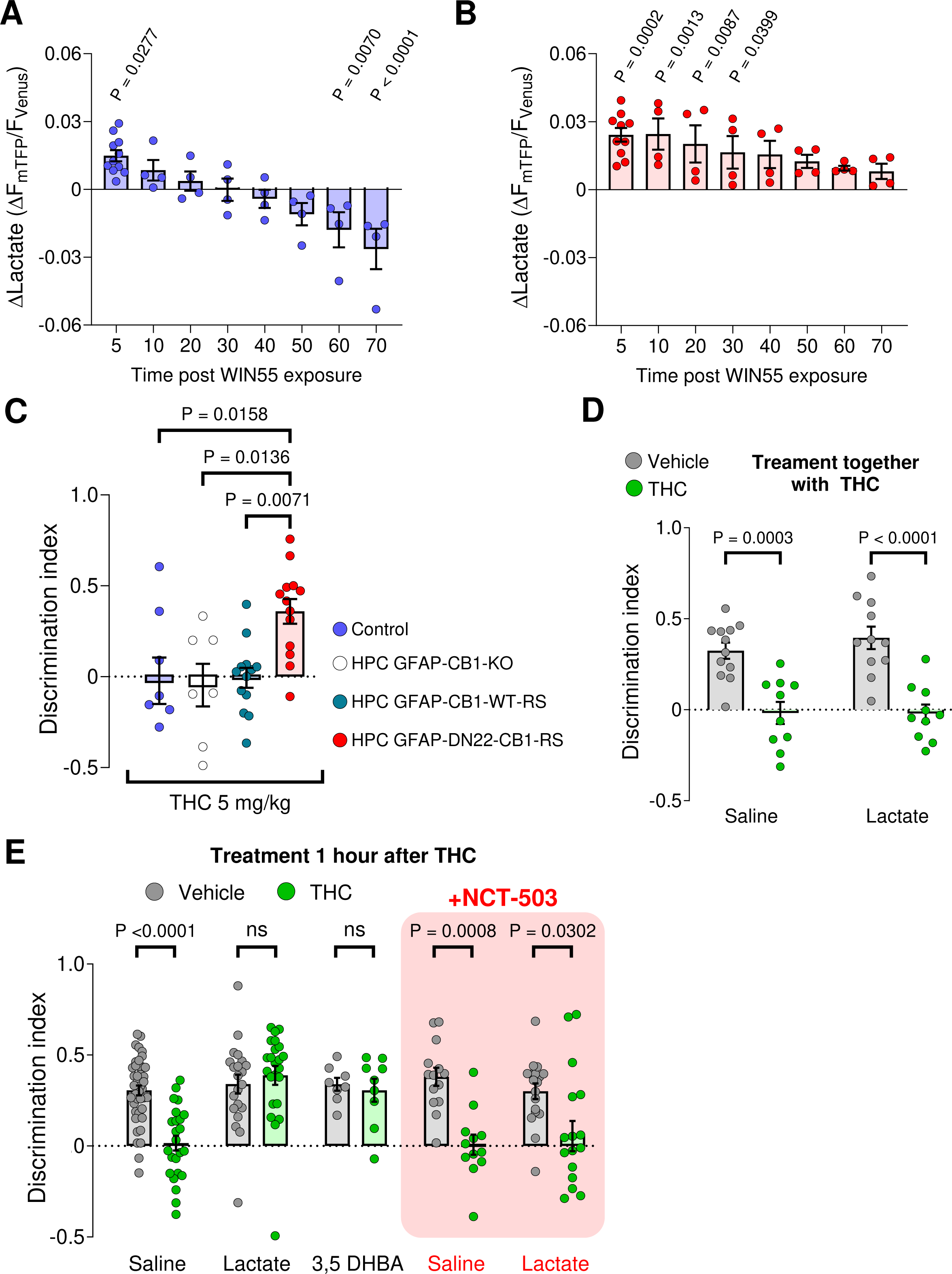
THC impairs NOR performance *via* mtCB1 receptor-dependent inhibition of the phosphorylated pathway. **A,** Intracellular lactate changes measured at different time points during a 70 min WIN55 exposure in CB1-WT astrocytes. N=4, 164 cells analyzed. Data from Fig. 1C (5 min quantification) is included in the graph. **B**, Intracellular lactate changes measured at different time points during a 70 min WIN55 exposure in DN22-CB1-KI astrocytes. N=4, 192 cells analyzed. Data from Fig. 1C (5 min quantification) is included in the graph. **C**, Summary of NOR performance in Control (blue), HPC-GFAP-CB1-KO (white), HPC-GFAP-CB1-RS (teal) and HPC-GFAP-DN22-CB1-RS (red) mice, treated with an IP injection of THC (5 mg/kg) immediately after acquisition phase. N = 7-13 mice per condition. **D**, NOR performance of mice treated either with an IP injection of either vehicle (grey) + saline, THC (5 mg/kg, green) + saline, vehicle + lactate (1 g/kg) or THC (5 mg/kg) + lactate (1 g/kg), immediately after the acquisition phase. N = 10 – 12 mice per condition. **E**, NOR performance of mice treated either with an I.P. injection of vehicle (grey) or THC (5 mg/kg, green) immediately after the acquisition phase. After 1-hour post THC or vehicle treatment, mice were treated with an IP injection of either saline, lactate (1g/kg), 3,5-DHBA (240 mg/kg), NCT-503 (6 mg/kg) + saline or NCT-503 + lactate. N = 8 – 49 mice per condition. Bars correspond to experiments average (mean ± SEM) and circles represent an experiment average (**A**,**B**) or represent individual animals (**C**,**D**,**E**). Statistical analysis was performed using a Mixed-effects model followed by Dunnett’s multiple comparison test (**A**,**B**) and a two-way ANOVA test by a Tukey’s multiple comparison test (**C**,**D**,**E**). See Supplementary Table 2 for more details.

We next asked whether this increase/decrease switch effect of cannabinoids might bear behavioral relevance. Using the same strategy as described in Figure 2, we found that the NOR-impairing effect of the plant-derived CB1 receptor agonist THC is absent in mice lacking mtCB1 receptors (HPC-GFAP-DN22-CB1-RS; Supplementary Table 1; Fig. 5C and Extended data Fig. 8C). This indicates a first mechanistic difference between physiological control of NOR performance by the endocannabinoid signaling (mtCB1 receptor-*independent*, Fig. 2B) and the disrupting effect of exogenous cannabinoids on the same process (mtCB1 receptor-*dependent*, Fig. 5C and Extended data Fig. 8C). Thus, we next wondered if this effect of THC might be linked to a mtCB1 receptor-dependent decrease of lactate. To address this issue, we tested whether lactate could rescue the effect of THC, similarly as it does in mice lacking CB1 receptors from astrocytes. The simultaneous administration of lactate and THC immediately after the acquisition of NOR did not alter the effect of the cannabinoid drug (Fig. 5D). Intriguingly, however, the administration of lactate 1 hour after THC completely reverted the impairment of long-term NOR performance (Fig. 5E and Extended Data Fig. 8D). Importantly, this effect of lactate was mimicked by 3,5-DHBA (Fig. 5E and Extended Data Fig. 8D), strongly suggesting that the effects of THC on NOR are due to a mtCB1 receptor-dependent decrease of lactate signaling through the HCAR1 receptor. This suggests that the rescuing effect of lactate on the THC effect might involve the PP cascade, similarly to physiological NOR memory consolidation. To address this possibility, we administered the PP inhibitor NCT-503 together with lactate 1 hour after THC treatment. Strikingly, this treatment abolished the rescuing effect of lactate (Fig. 5E and Extended Data Fig. 8D). Altogether, these data clearly indicate that lactate signaling is required to consolidate NOR memory. This signaling is disrupted by the insufficient production of lactate occurring both in absence of physiological CB1 receptor activity and during pharmacological stimulation of mtCB1 receptors in astrocytes.

## DISCUSSION

This study reveals a novel astrocyte-dependent metabolic interaction between lactate signaling, the phosphorylated pathway, synaptic D-serine availability and cognitive performance in mice. Our data are compatible with a scheme whereupon endogenous activation of astroglial CB1 receptors leads to a temporary increase of lactate, which in turn activates HCAR1 signaling, switching glycolysis towards serine production and ultimately providing the NMDAR activity necessary for physiological novel object recognition (Extended Data Fig. 9). Strikingly, the prolonged exposure of astrocytes to exogenous cannabinoids results in impaired cognitive performance through a specular mechanism that involves activation of astroglial mtCB1 receptors and inhibition of the same lactate signaling (Extended Data Fig. 9). Thus, these data provide a novel mechanistic link between astroglial energy metabolism and gliotransmission, and they explain the differential effects of physiological *versus* pharmacological impact of CB1 receptor activation on cognitive processes.

Astroglial CB1 receptors have been shown to control synaptic plasticity in different brain regions and to determine cognitive processes^12, 14, 47–49^. The regulation of astrocyte calcium signaling is generally indicated as the cellular mechanism underlying these functions^43, 50, 51^. Only recently, cannabinoid signaling was linked to specific metabolic control of behavior, showing that persistent activation of astroglial mtCB1 receptors can reduce lactate production by astrocytes and induce a bioenergetic crisis in neurons, eventually resulting in impairment of social interactions^16^. Here, using lactate-sensitive fluorescent biosensors, we confirmed that long-term application of cannabinoids reduces lactate levels in astrocytes^16^, but we found also that short-term activation of astroglial CB1 receptors stimulates lactate production and release. Consistent with this observation, the exogenous administration of lactate is sufficient to rescue the NOR impairment of GFAP-CB1-KO mice. Lactate levels increase in the brain parenchyma during neural workload^34^, a phenomenon explained by a shift from complete to partial glucose oxidation and known as aerobic glycolysis^52^. Several signals have been proposed to initiate this metabolic shift, such as glutamate^53, 54^, extracellular K^+^ rises^55, 56^ and others^24, 57, 58^. The present data indicate that astroglial CB1 receptor signaling participates in these processes, suggesting that it can trigger aerobic glycolysis and concomitant increase in extracellular lactate levels. When compared to other brain signals, the effect of CB1 receptor activation on lactate metabolism share a similar time scale to the effects of extracellular K^+^ rises, which activate astrocyte glycolysis within seconds^59, 60^ and promote a phenomenon of metabolic recruitment of neighboring astrocytes^56^. However, endocannabinoid signaling is thought to be highly local^1, 45, 46^, a characteristic similar to glutamate which does not diffuse far from its release sites^61^. This suggests that CB1 receptors may provide fast and local signaling to trigger astrocyte lactate metabolism, working in parallel or synergistically with other brain signals that control astrocyte metabolic functions.

Our data show that PKC activity is required for this phenomenon. PKC signaling is very complex and formed by several isozymes exerting different control of cellular activity^62–64^. Astrocytes express different types of PKC isozymes^65^, but their role in astrocyte metabolic function is almost unexplored^66, 67^. However, in other cell types PKC signaling modulates glucose uptake^68–73^, glycolytic flux^74–77^ and lactate dehydrogenase activity^78^. Thus, it will be very interesting to address the detailed characterization of the intracellular machinery linking CB1 receptors, PKC and lactate metabolism. In this context, the present data demonstrate that deletion of (mt)CB1 receptors does not alter basal lactate metabolism, and amplitude and kinetics of lactate accumulation induced by OXPHOS inhibition. However, astroglial lactate metabolism is regulated by diverse stimuli, including neuronal signals^22, 56^, cellular stress or alterations in mitochondrial respiration^17, 79^. These stimuli can trigger a plethora of intracellular cascades, which could each interact with (mt)CB1 receptors. Thus, we cannot presently exclude that potential alterations in different mechanisms regulating lactate production might exist in cells lacking (mt)CB1 receptors. Further studies will deal with this potential issue by exploring how CB1 receptors might interact with other stimuli that modulate astroglial lactate metabolism.

Noteworthy, our present and previous^16^ data altogether indicate that activation of CB1 receptors results in a biphasic time-dependent modulation of astrocyte lactate metabolism. We previously showed that persistent pharmacological activation (24 hours) of astroglial mtCB1 receptors decreases lactate production, thereby causing neuronal bioenergetic stress and impairing social interactions in mice^16^. Conversely, in the present study, we report that a short-term activation (< 10 min) of non-mitochondrial astrocyte CB1 receptors results in a transient increase of intracellular lactate. Here we found that this switch between increase and decrease of lactate levels by cannabinoids occurs within one hour, indicating that time plays a fundamental role in determining the modalities (e.g., functional engagement of mitochondrial or non-mitochondrial CB1 receptors) and the outcome of cannabinoid signaling. This bimodal and subcellular-specific action on lactate levels resembles the recently described differential involvement of neuronal plasma membrane and mitochondrial CB1 receptors in cannabinoid-induced antinociception and catalepsy, respectively^11^. However, those subcellular-specific effects of cannabinoids occur simultaneously, whereas the differential impact on lactate levels involve an important temporal component. In this context, it is interesting to note that the physiological control of NOR performance and its pharmacological impairment by cannabinoid agonists intersect at the same molecular pathway, which is promoted or inhibited in a time-dependent manner. Indeed, the physiological activation of non-mitochondrial CB1 receptors seems to transiently increase lactate levels and the successive signaling, whereas persistent presence of CB1 agonists engages mtCB1 receptors to decrease the metabolite levels. Our data show that this temporal dichotomy bears important functional and behavioral consequences. Indeed, they are consistent with a scenario whereupon physiological and pharmacological activation of CB1 receptors trigger opposite effects mediated by different subcellular populations of the receptor (Extended Data Fig. 9). Thus, endocannabinoids mobilized during physiological consolidation of NOR memory activate non-mitochondrial CB1 receptors to increase lactate levels, HCAR1 signaling, PP activity, serine availability and NMDAR synaptic functions (Extended Data Fig. 9). Conversely, pharmacological administration of exogenous cannabinoid agonists reaches astroglial mtCB1 receptors, triggering the exact opposite cascade, ultimately impairing NOR memory consolidation (Extended Data Fig. 9).

Lactate has traversed a long way from being initially viewed as little more than mere cellular waste, to become a relevant metabolite for brain physiology^22, 34^. Astrocyte-derived lactate has been shown to modulate neuronal functions *via* multiple mechanisms^80–90^. Most of these proposed mechanisms, however, are linked to the consumption of lactate as an energy substrate for neuronal activity, which likely fuel mitochondrial energy production, but also increases the cytosolic NADH levels, which potentiate NMDAR activity and promote expression of plasticity genes^84^. Indeed, the potentiation of NMDAR activity by lactate in our electrophysiological experiments is significantly blunted by blockade of the phosphorylated pathway by NCT-503. However, a non-significant “residual” level of potentiation was consistently observed in these experiments, suggesting that multiple mechanisms likely link lactate to synaptic functions. This said, the present data reveal a receptor-dependent impact of lactate on cognitive processes: the control of D-serine synthesis and its synaptic availability. This process requires the HCAR1 signaling-dependent activation of the phosphorylated pathway in astrocytes^38, 41^. This novel signaling process likely works in parallel with the other described effects of lactate in the brain that have been shown to be essential to sustain learning and memory processes (including monocarboxylate transporter-dependent uptake and metabolism)^84, 89–92^. It is reasonable to speculate that these multiple mechanisms might allow astrocytes minimizing the resources required to cooperate with neurons to fulfill different behavioral tasks.

Important to note, activation of HCAR1 has been suggested to reduce neuronal functions^87, 88, 93^, which might contradict our results. However, careful dose dependent experiments using the 3,5-DHBA agonist in hippocampal slices showed that HCAR1 activation exerts a biphasic effect on CA1 pyramidal neurons, with low doses decreasing and high dose increasing their excitability^85^. The concentration of 3,5-DHBA (1 mM) used here corresponds to a high dose that increases CA1 pyramidal cells excitability^85^. Thus, our results are in line with the literature and support a more complex view of HCAR1 signaling in the brain. On the other hand, the molecular link(s) between HCAR1 signaling and the phosphorylated pathway are currently unknown, and future studies will address the details of this novel signaling cascade, investigating the localization of the involved HCAR1 (e.g. neurons or astrocytes)^94–96^ and the molecular components of the involved signaling pathway. For instance, HCAR1 are generally coupled to Gi proteins^35, 36^. Although virtually nothing is known on the molecular regulation of the enzymes of the phosphorylated pathway, it is possible that G protein-triggered signaling might impact the activity of at least some of them (e.g. *via* specific kinase- and/or calcium-dependent processes).

The spatio-temporal pattern of lactate-stimulated L-serine activity might also play a potentially important mechanistic role in the link between CB1 receptors and NMDAR activity. It is possible that the CB1-dependent increase of lactate and the production of L-serine coexist in a single astrocyte, but they are separated in time. In other words, upon CB1 receptor activation, astrocytes may first accumulate lactate, and then trigger HCAR1 signaling to promote L-serine. Alternatively, CB1 receptor-stimulated astrocytes might act as net lactate providers, whereas another astrocyte population expressing HCAR1 may possibly be responsible for the L-serine production upon stimulation by lactate. This possibility might be similar to the specific gating of striatal circuits by distinctive astrocyte subpopulations^97^, and the recently proposed secondary metabolic recruitment induced by the diffusion of lactate away from its release site^56^. Further focused studies will be necessary to fully unveil the mechanisms explaining the interaction between lactate, HCAR1 signaling and L-serine. However, the discovery of the existence of such interaction reveals key connections between metabolic processes (lactate) and synaptic signaling (D-serine), which are required for cognitive processes.

D-serine is a gliotransmitter and co-agonist of synaptic NMDARs^14, 41, 98–100^. Despite that both neurons and astrocytes can likely release this amino acid^39, 49^, astrocytes are the main producers of its precursor L-serine^40^, thereby representing the main controllers of the total amount of D-serine in the brain. By showing that astrocyte-borne lactate promotes D-serine signaling *via* HCAR1-dependent stimulation of the phosphorylated pathway, our results underscore another way by which astrocytes modulate brain functions. Thus, this study contributes to a unifying concept of astrocyte metabolic functions and gliotransmission. These two processes are often considered as independent entities. For instance, researchers either study just astrocytes metabolic processes or the impact of astrocytes functions on synaptic activity^101–104^, with little interactions between these points of view. This is particularly true when the roles of astrocytes are investigated in the frame of cognition and high mental processes. In this study, we merged the new observation that type-1 cannabinoid CB1 receptors can rapidly and transiently promote lactate metabolism and the regulation of D-serine synaptic functions by endocannabinoid signaling in astrocytes. Moreover, we identified the same metabolic/signaling cascade as the target of prolonged pharmacological impairment of cognition by cannabinoid drugs. Therefore, these data show the tight functional link existing between metabolic and signaling processes. This unified notion will have to be taken into account in future studies aimed at understanding the role of astrocytes in promoting and controlling brain functions and behavior, and at investigating neurological and psychiatric disorders characterized by cognitive impairment.

**Table 1.** Details of the double-viral rescue approach to study mtCB1 receptor involvement in NOR performance.

| Mice | AAV-1* | AAV-2* | Outcome | Mitochondrial Localization of CB1? | Name of used mutant mice<sup>2</sup> |
| --- | --- | --- | --- | --- | --- |
| CB1-flox | GFAP-GFP | CAG-DIO-Empty | CB1-WT | yes | Control |
| CB1-flox | GFAP-Cre | CAG-DIO-Empty | CB1-KO in GFAP positive cells | no | HPC-GFAP-CB1-KO |
| CB1-flox | GFAP-Cre | CAG-DIO-CB1-GFP | Re-expression of CB1-WT in GFAP positive cells (CB1 in all subcellular locations) | yes | HPC-GFAP-CB1-RS |
| CB1-flox | GFAP-Cre | CAG-DIO-DN22-CB1-GFP | Re-expression of DN22-CB1 in GFAP positive cells (CB1 excluded from mitochondria) | no | HPC-GFAP-DN22-CB1-RS |

\*Injected into the hippocampus (HPC)

## ACKNOWLEDGEMENTS

We would like to thank Delphine Gonzales, Nathalie Aubailly, Ruby Racunica, Jean-Baptiste Bernard and all the personnel of the Animal Facilities of the NeuroCentre Magendie for mouse care. We also thank the genotyping platform of the Neurocentre Magendie for the help in the experiments. The microscopy was done in the Bordeaux Imaging Center a service unit of the CNRS-INSERM and Bordeaux University, member of the national infrastructure France BioImaging supported by the French National Research Agency (ANR-10-INBS-04). We thank all past and present members of Marsicano’s lab for useful discussions and for their invaluable support. This study was funded by Inserm (to G.M., A.P. and S.H.R.O.); CNRS (to A.P. and S.H.R.O.); the European Research Council (Micabra, ERC-2017-AdG-786467, to G.M.); Fondation pour la Recherche Medicale (DRM20101220445, to G.M); EMBO Long-term Fellowship ALTF87-2018 (to I.F-M.); the Human Frontiers Science Program (to G.M.); Region Aquitaine (CanBrain, AAP2022A-2021-16763610 and -17219710 to A.-K.B.-S., L.P., A.P. and G.M.); French State/Agence Nationale de la Recherche (ERA-Net Neuron CanShank, ANR-21-NEU2-0001-04, to G.M), (CaMeLS, to G.M., A.P. and G.B.),(Hippobese, ANR-23-CE14-0004-03, to G.F. and G.M.), (BrainFuel, ANR-21-CE44-0023-01 to A.-K.B.-S., L.P and AP.), (Excigly, ANR-20-CE16-0009-03 to S.H.R.O.), (Astrocom ANR-19-CE16-0015 to A.P.), (ANR-19-CE14-0039 to L.B.); the French government in the framework of the University of Bordeaux’s IdEx “Investments for the Future” program / GPR BRAIN_2030, the Japan Society for the Promotion of Science (21K14738, to Y.N. and 19H05633, to R.E.C.) and the Japan Science and Technology Agency (JPMJPR22E9, to Y.N.), Fondation Alzheimer (to G.B.), the NextGenerationEU/PRTR and Agencia Estatal de Investigación (10.13039/501100011033; PID2019-105699RB-I00; PID2022-138813OB-I00 and PDC2021-121013-I00 to J.P.B); European Comission action HORIZON-TMA-MSCA-DN (ETERNITY, 101072759, to J.P.B), La Caixa Research Health grant HR23-00793 (to J.P.B. and G.M.) and Fondecyt 1230145 & BMBF-ANID 180045 (to L.F.B.).

## AUTHORS CONTRIBUTION

IFM conceived the study, wrote the manuscript and performed most experiments; GL, UBF, NB and SM performed behavioral and electrophysiological experiments; PH, TDT, RS, LB, AC and BFM helped with experiments; FJK and DG performed the histology analyses; YN and REC generated and provided sensor constructs; FD, CC, GF, AKBS, LP, JPB, GB, LFB and SHRO provided theoretical support and ideas; AP conceived the study; GM conceived the study and wrote the manuscript; all authors edited and approved the manuscript.

## Extended Data Figures legends

**Extended Data Figure 1.**
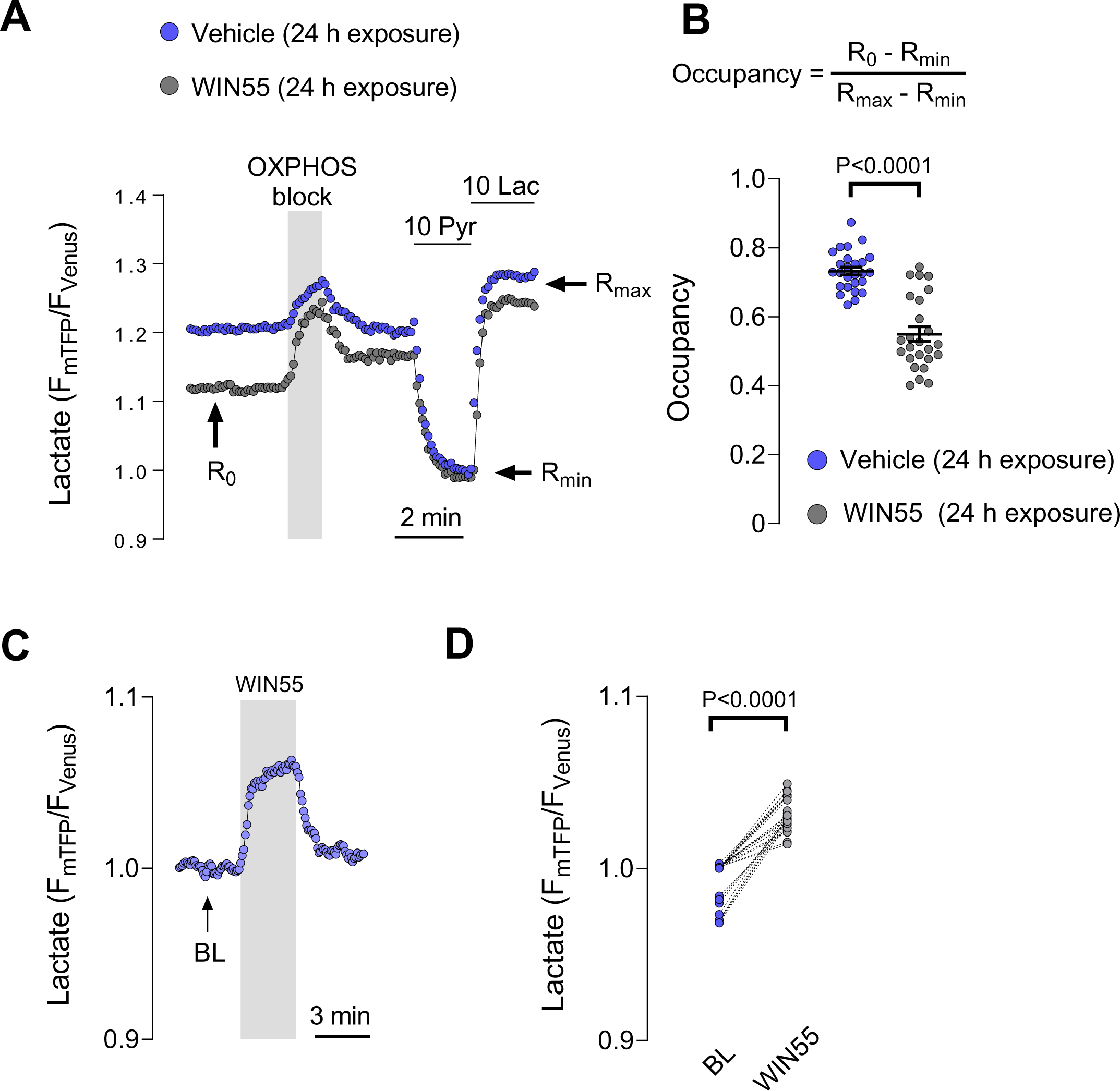
Differential effect of cannabinoids on astrocyte lactate level. **A**, Intracellular lactate imaging in astrocytes previously incubated with WIN55 (2 μM) or vehicle (DMSO) for 24 hours. After treatment, and to determine the basal lactate level (occupancy), cells were imaged and exposed sequentially to an OXPHOS blocker (5 mM sodium azide), pyruvate (10 mM) and lactate (10 mM). R_0_, basal ratio. R_min_, minimum ratio. R_max_, maximum ratio. Data was normalized to R_min_ to emphasize the difference in R_0_. **B**, Basal lactate level (occupancy) after 24 hours treatment with WIN55 (2 μM) or vehicle (DMSO). Data was computed as occupancy = (R_0_-R_min_)/(R_max_-_Rmin_), using R_0_, R_min_ and R_max_ from experiments similar to panel A. Vehicle, n=3, 26 cells. WIN55, n=3, 25 cells. **C**, Intracellular lactate imaging in astrocytes acutely exposed to WIN55 (2 μM). BL = baseline. **D**, Summary of intracellular lactate level at baseline (BL) and after 3 min exposure to WIN55 (2 μM), in experiments similar to Extended Data Fig. 1C, n=3, 26 cells. Data correspond to representative cells (**A**,**C**). Circles in scatter and before-after plots correspond to single cells (**B**,**D**). Statistical analysis was performed using a two-tailed unpaired t-test (**B**) and two-tailed paired t-test (**D**). See Supplementary Table 3 for more details.

**Extended Data Figure 2.**
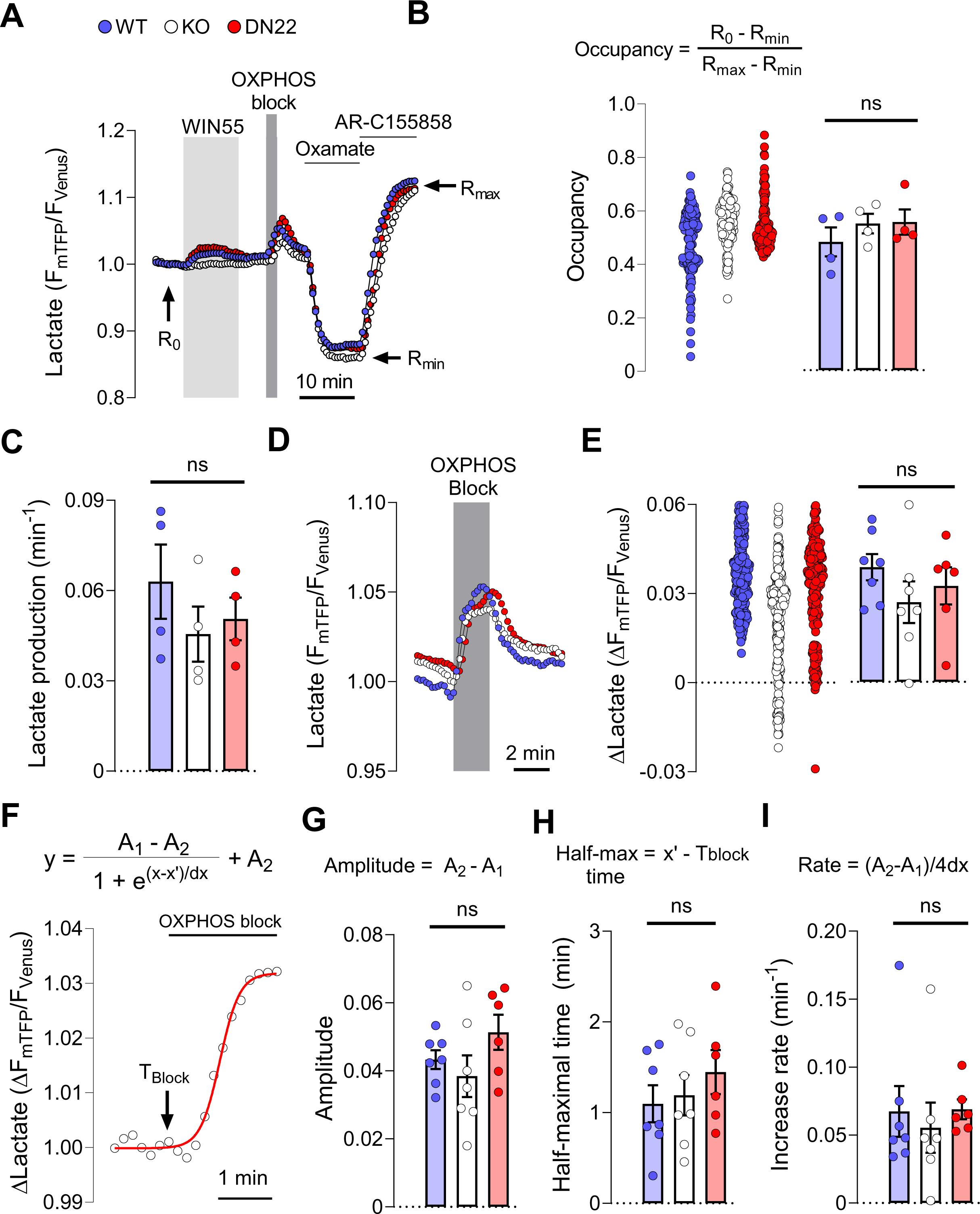
The basal lactate level and accumulation upon mitochondria inhibition is not altered by CB1 receptor subcellular localization. **A**, Intracellular lactate imaging in astrocytes. To determine the basal lactate level (occupancy), cells exposed sequentially to WIN55 (1 μM), OXPHOS block (5 mM azide), Oxamate (6 mM) and AR-C155858 (1 μM). R_0_, basal ratio. R_min_, minimum ratio. R_max_, maximum ratio. Average of 4 independent experiments. Cells: WT=158, KO=145, DN22=154. **B**, Basal lactate level (occupancy) in CB1-WT, CB1-KO and DN22-CB1-KI astrocytes. Data was computed as occupancy = (R_0_-R_min_)/(R_max_-_Rmin_), using R_0_, R_min_ and R_max_ obtained from experiments similar to panel A. N = 4, cells analyzed: CB1-WT=158, CB1-KO=145, DN22-CB1-KI =154. **D**, Intracellular lactate accumulation induced by OXPHOS block (5 mM sodium azide). Average of several cells in a representative experiment (CB1-WT=32, CB1-KO=48, DN22-CB1-KI =36 cells). **E**, Summary of intracellular lactate levels after 2 min of OXPHOS block (5 mM sodium azide), in experiments similar to those shown in panel C. CB1-WT: n=7, 247 cells. CB1-KO: n=7, 227 cells. DN22-CB1-KI: n=6, 205 cells. **F**, Representative non-linear fitting of a sigmoidal model (Boltzmann equation, on top) to the lactate increase induced by OXPHOS blocking. The fitted parameters A_1_, A_2_, x’ and dx were used to compute the amplitude, half-maximal time and increase rate presented in panel G-I. T_block_, time of exposure to OXPHOS blocker sodium azide. **G**, Amplitude of lactate changes induced by OXPHOS block obtained from a non-linear fitting data. CB1-WT: n=7; CB1-KO: n=7; DN22-CB1-KI: n=6. **H**, Half-maximal time of lactate changes induced by OXPHOS block obtained from a non-linear fitting data. CB1-WT: n=7; CB1-KO: n=7; DN22-CB1-KI: n=6. **I**, Half-maximal time of lactate changes induced by OXPHOS block obtained from a non-linear fitting data. CB1-WT: n=7; CB1-KO: n=7; DN22-CB1-KI: n=6. Data corresponds to the experiments average and represented as mean±SEM (**A**,**D**). Circles in scatter plots correspond to single cells (**B**,**E**). Bars correspond to experiments average (mean±SEM) and circles represent individual experiment average (**B**,**C**,**E**,**G**,**H**,**I**). Statistical analysis was performed using Kruskal-Wallis test followed by Dunn’s multiple comparison test (**B**), One-way ANOVA followed by Tukey’s multiple comparison test (**C**,**E**,**G**,**H**,**I**). See Supplementary Table 3 for more details.

**Extended Data Figure 3.**
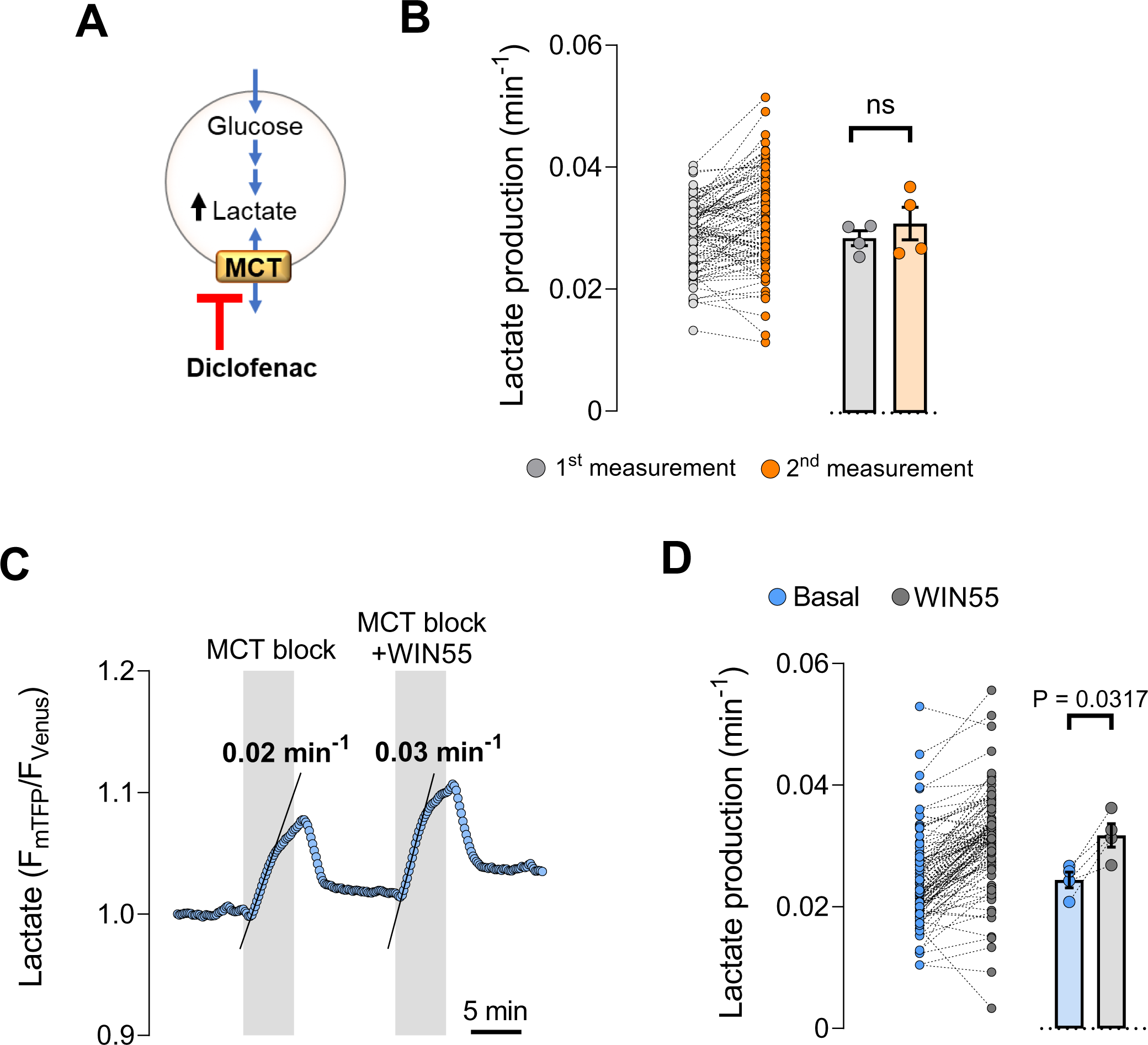
Activation of astroglial CB1 receptors increases lactate production. **A**, Transport stop protocol for measurement of lactate production. Diclofenac is a broad inhibitor of monocarboxylate transporter (MCT) activity. The blockade of MCT causes an intracellular lactate accumulation that is proportional to its rate of production. **B**, Summary of two sequential measurements of basal lactate production with diclofenac. N=4, 95 cells analyzed. **C**, Measurement of lactate production before and during exposure to WIN55 (1 μM). The production rate is indicated with a solid line above the corresponding lactate accumulation. **D**, Summary of the lactate production rates before (pale blue circles) and during exposure to WIN55 (grey circles), computed from experiments similar to panel E. N=4, 102 cells. Data corresponds to representative cell I. Circles in before – after plots correspond to single cells (**B**,**D**). Bars correspond to experiments average (mean±SEM) and circles represent individual experiment average (**B**,**D**). Statistical analysis was performed using a paired two-tailed paired t-test (**B,D**). See Supplementary Table 3 for more details.

**Extended Data Figure 4.**
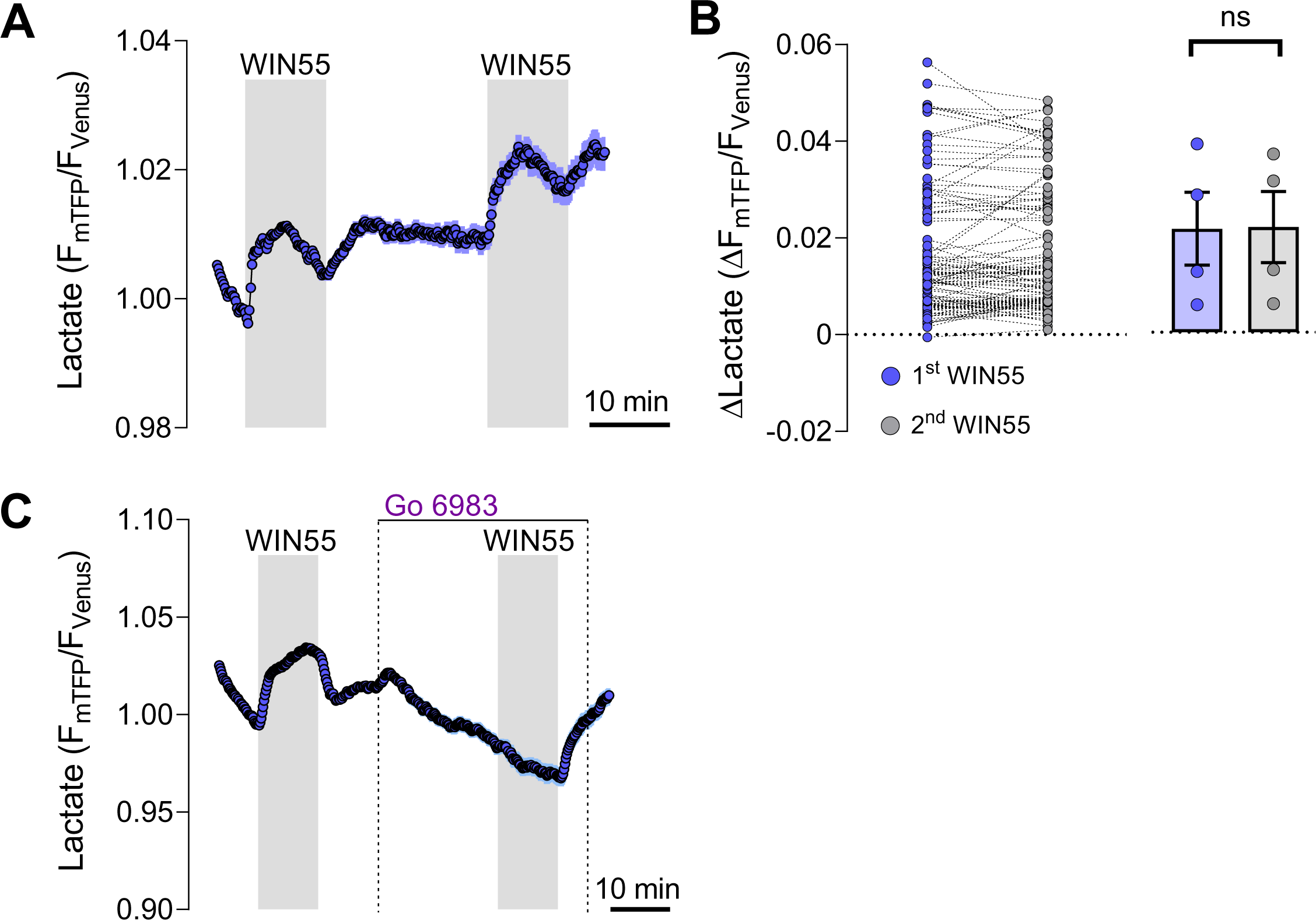
The PKC signaling controls the CB1 receptor-mediated intracellular lactate increase. **A,** Intracellular lactate measurement during exposure two sequential exposure to WIN55 (1 μM). N=1, 42 cells. **B**, Quantification of lactate change induced by the first (blue) and second (grey) WIN55 exposure. N=4, 114 cells analyzed. **C**, Intracellular lactate measurement during the sequential exposure to WIN55 (1 µM), Go 6983 (5 µM) and WIN55 + Go 6983. N=1, 40 cells. Data corresponds to the average of a single experiment (**A**,**D**). Circles in before – after plots correspond to single cells (**B**). Bars correspond to experiments average (mean±SEM) and circles represent individual experiment average (**B**). Statistical analysis was performed using a paired two-tailed paired t-test (**B**). See Supplementary Table 3 for more details.

**Extended Data Figure 5.**
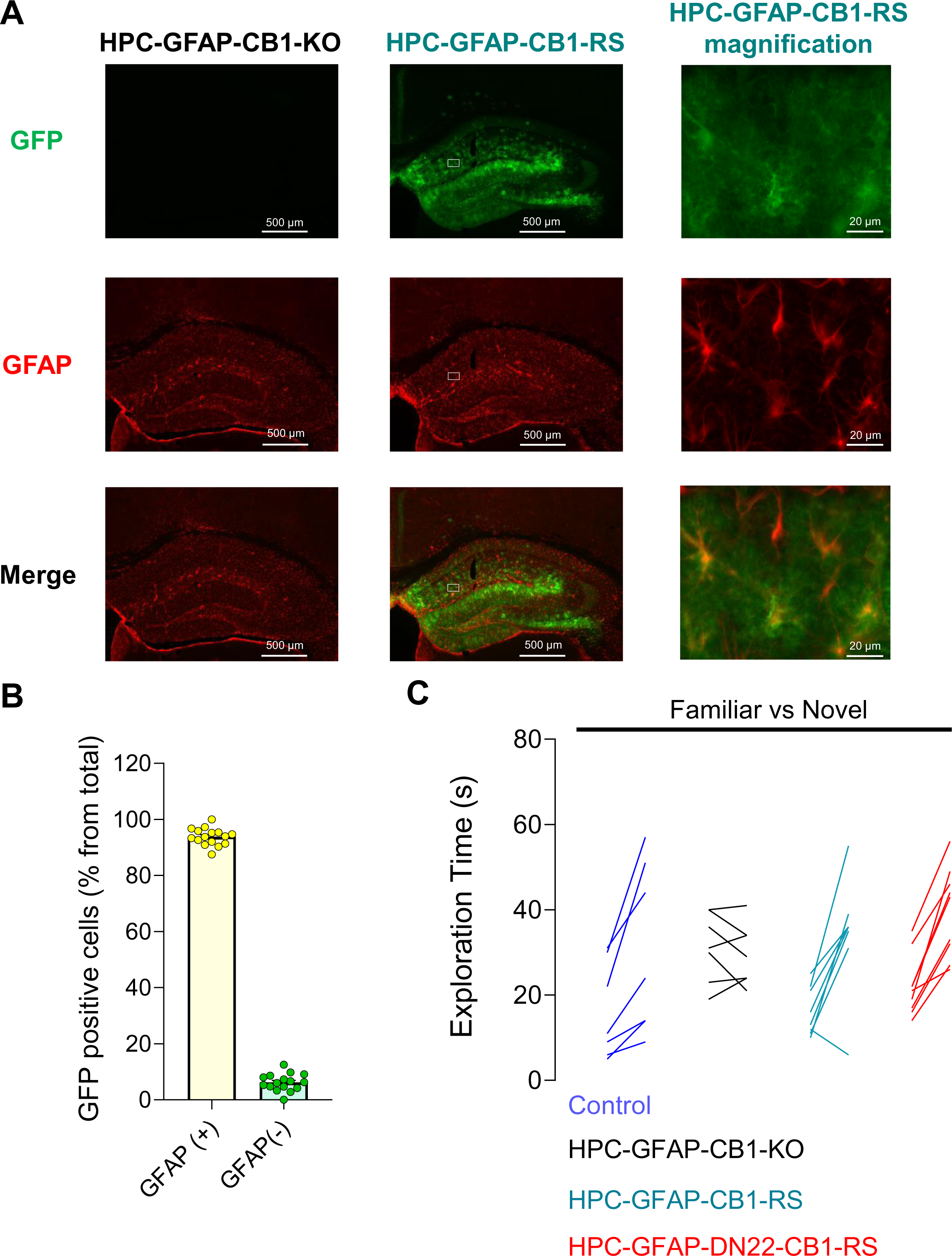
The mitochondrial localization of CB1 receptors is not necessary for physiological novel object exploration. **A,** Histological analysis of the expression of CB1-GFP and the astrocyte marker GFAP, in hippocampus sections obtained from HPC-GFAP-CB1-KO and HPC-GFAP-CB1-WT-RS. The white boxes inside the HPC-GFAP-CB1-WT-RS images correspond to the magnification site shown in the third column of images. **B**, Quantification of GFP-positive cells in the CA1 region of the hippocampus of HPC-GFAP-CB1-WT-RS mice. N = 4 mice, 30 – 91 cells analyzed in 4 sections per mice. **C**, Exploration times of familiar versus novel objects in the NOR task, from Control (blue lines), HPC-GFAP-CB1-KO (black lines), HPC-GFAP-CB1-WT-RS (teal lines) and HPC-GFAP-DN22-CB1-RS (red lines) animals, n = 7-9 mice per condition. A single line corresponds to an individual animal. See Supplementary Table 3 for more details.

**Extended Data Figure 6.**
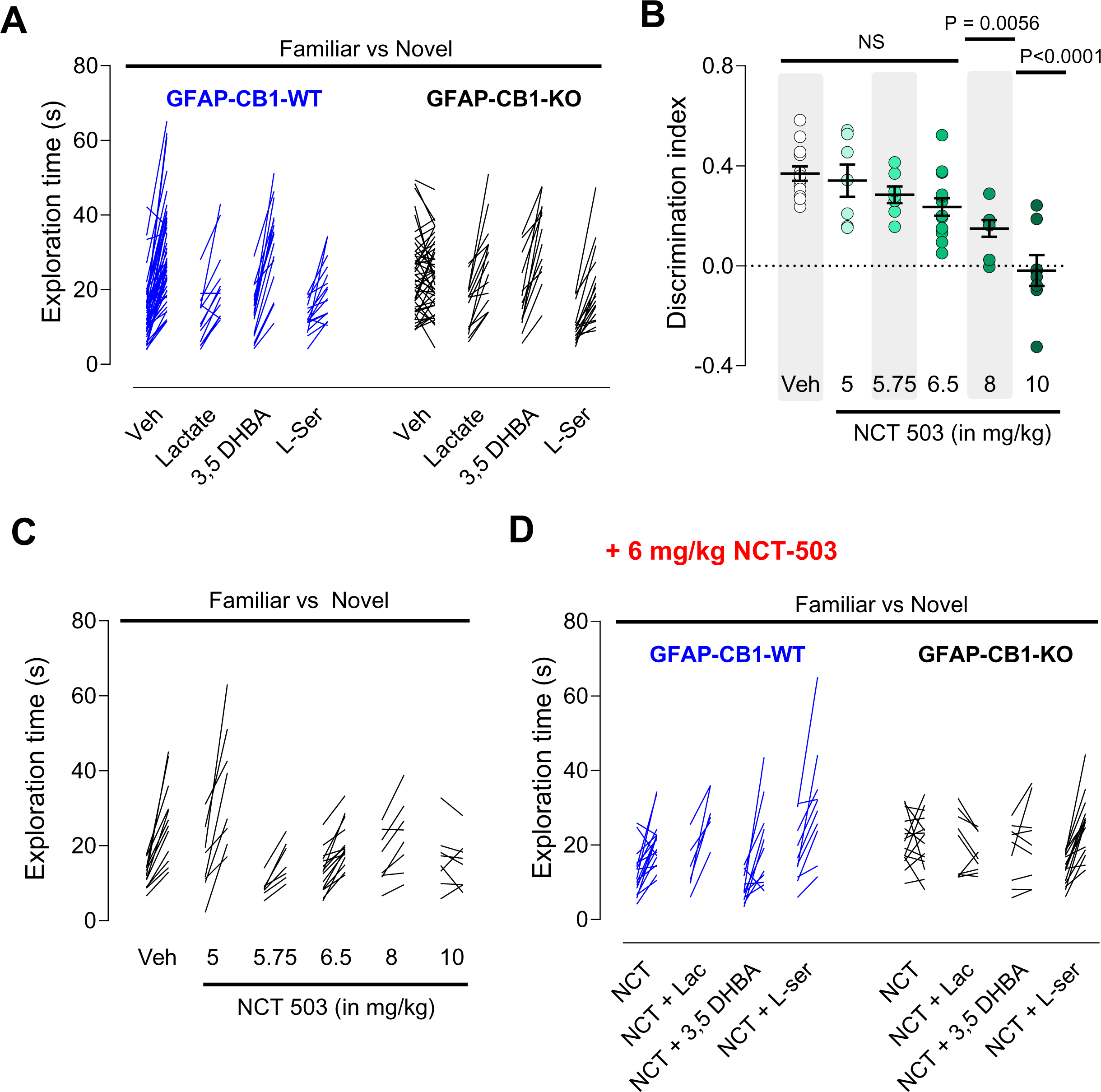
Inhibition of the phosphorylated pathway impairs long-term NOR memory in WT mice and in lactate-treated GFAP-CB1-KO mice. **A**, Exploration time of familiar versus novel objects in the NOR task of GFAP-CB1-WT (blue lines) and GFAP-CB1-KO mice (black lines) mice, treated either with vehicle (veh), 1 g/kg lactate (Lac) or 0.5 g/kg L-serine (L-Ser), immediately after the acquisition phase. GFAP-CB1-WT, n=13-18 animals. GFAP-CB1-KO, n= 15-23 animals. **B**, NOR performance in wild-type mice treated either with vehicle or incremental doses of NCT-503. N=7-15 mice per condition. **C**, Exploration time of familiar versus novel object in the NOR task of wild-type mice treated either with vehicle or incremental doses of NCT-503. N= 7-15 mice per condition. **D**, Exploration time of familiar versus novel object in the NOR task, of mice treated either with vehicle + 6 mg/kg NCT-503 (NCT), 1 g/kg lactate + 6 mg/kg NCT-503 (NCT + Lac) or 0.5 g/kg L-serine + 6 mg/kg NCT-503 (NCT + L-Ser), immediately after the acquisition phase. GFAP-CB1-WT mice (blue lines), n=6-12 animals. GFAP-CB1-KO mice (black lines), n= 9-16 animals. A single line corresponds to an individual animal (**A**, **C**, **D**). Data is presented as scatter plot with the line and whisker corresponding to the mean±SEM and circles to individual animals (**B**). Statistical analysis was performed with a One-way ANOVA followed by Tukey’s multiple comparison test (**B**). See Supplementary Table 3 for more details.

**Extended Data Figure 7.**
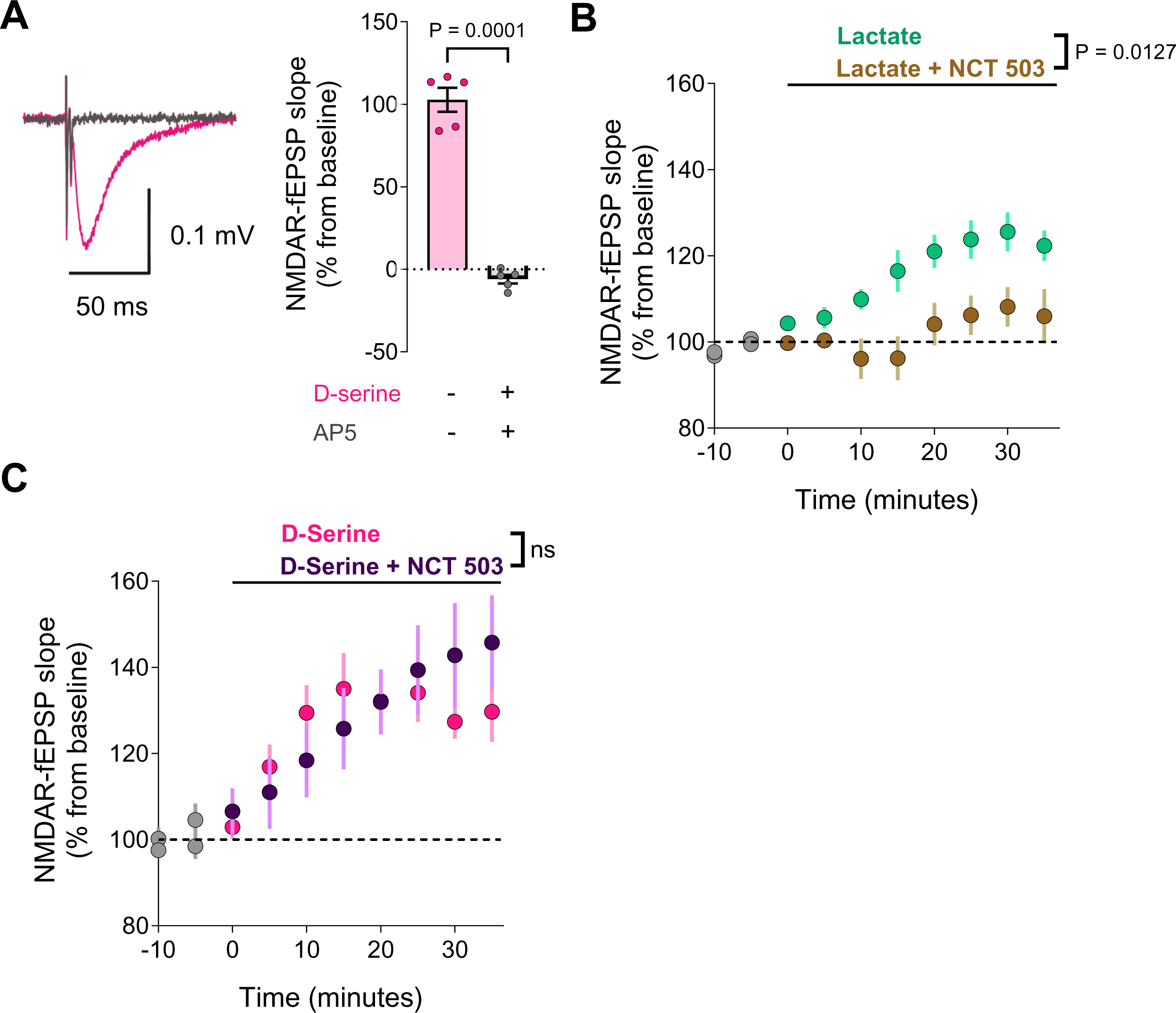
Lactate requires the phosphorylated pathway to potentiate NMDAR function. **A**, Representative averaged traces from 20 consecutives sweeps evoked in the presence of, before (in magenta) and after bath application of D-AP5 (50 µM) + D-serine (50 µM). Quantification of the NMDAR-fEPSP slopes in presence of D-serine (50 μM), before and after application D-AP5 (50 μM) are shown in the bar plot, n=5. **B**, NMDAR-mediated fEPSP slopes in the presence of lactate (data from Fig. 4A, n=9) and lactate + NCT-503 (data from Fig. 4D, n=6). **G**, NMDAR-fEPSP slopes induced by D-serine (same as Fig. 4A, n=6) and D-serine after NCT-503 preincubation (data from Fig. 4D, n=6). Bars correspond to experiments average (mean±SEM) and circles represent individual experiment average (**A**). Data corresponds to the experiments average and represented as mean±SEM. Data points were averaged every 5 mins (**B**,**C**). Statistical analysis was performed using a two-tailed paired t-test (**A**) and two-way ANOVA (**B**,**C**). See Supplementary Table 3 for more details.

**Extended Data Figure 8.**
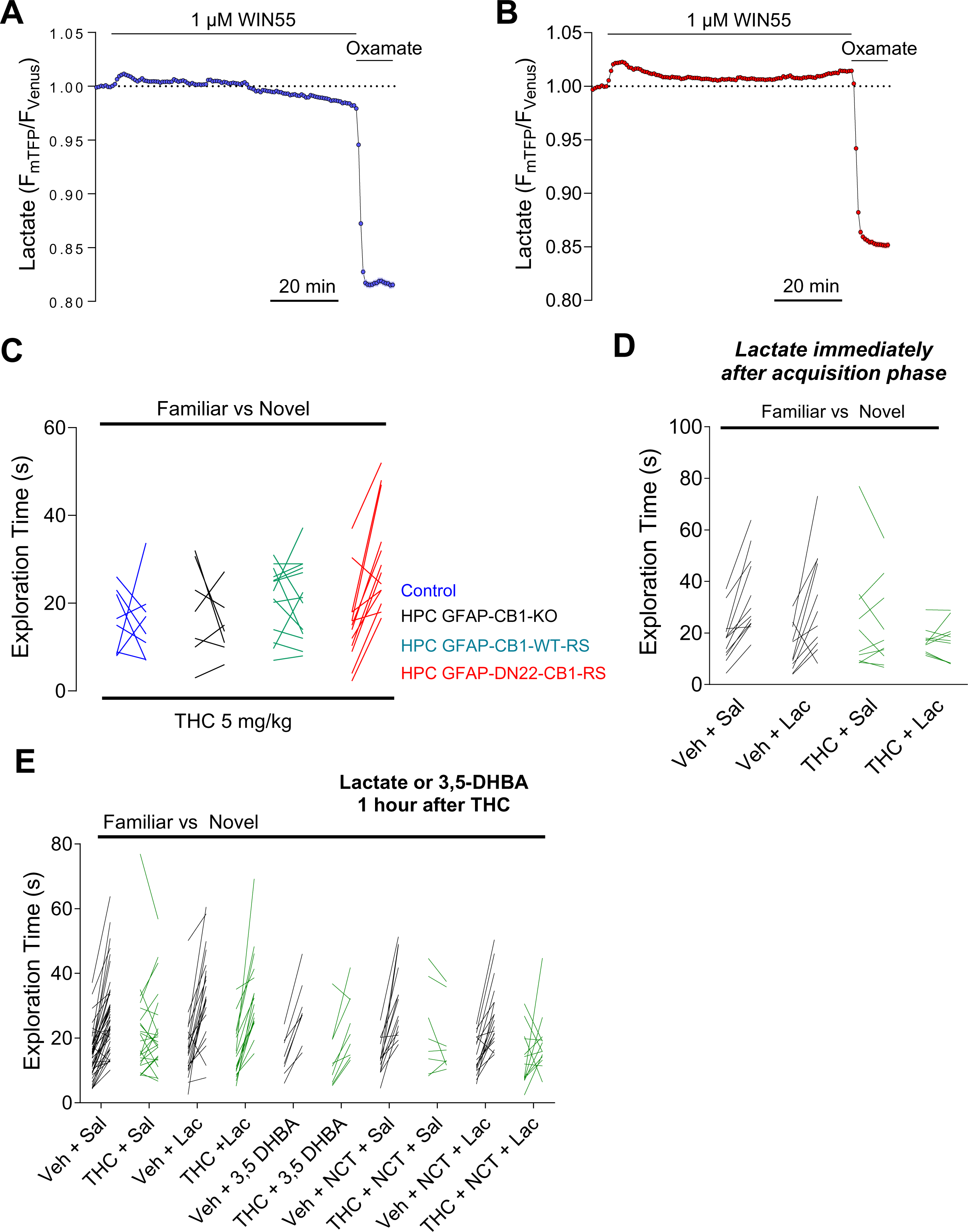
Lactate rescues the THC-mediated impairment in novel object exploration via HCAR1 signaling and L-serine production. **A**, Intracellular lactate imaging in CB1-WT astrocytes exposed to WIN55 (1 μM) during 70 min. After this, cells were exposed to oxamate to deplete lactate levels for biosensor calibration. **B**, Intracellular lactate imaging in DN22-CB1-KI astrocytes exposed to WIN55 (1 μM) during 70 min. After this, cells were exposed to oxamate to deplete lactate levels for biosensor calibration. **C**, Exploration times of familiar versus novel objects in the NOR task, from Control (blue lines), GFAP-CB1-KO (black lines), GFAP-CB1-WT-RS (teal lines) and GFAP-DN22-CB1-RS (red lines) animals treated with THC (5 mg/kg immediately after the acquisition phase. N = 7 – 13 mice per condition. **D**, Exploration time of familiar versus novel objects in the NOR task of mice treated with an IP injection of either vehicle (veh) + saline (sal), vehicle + lactate (lac, 1 g/kg), THC (5 mg/kg) + saline or THC (5 mg/kg) + lactate (1 g/kg), immediately after the acquisition phase. N= 10 – 12 mice per condition. **E**, Exploration time of familiar versus novel objects in the NOR task of mice treated with an IP injection of either vehicle (veh) or THC (5 mg/kg), immediately after the acquisition phase. After 1-hour post-THC treatment, mice were treated with an IP injection of either saline (sal), lactate (lac, 1g/kg), 3,5-DHBA (240 mg/kg), NCT-503 (NCT, 6 mg/kg) + saline or NCT-503 + lactate. N = 8 – 49 mice per condition. Experiments correspond to a representative cell (**A**, **B**). A single line corresponds to an individual animal (**C**, **D**, **E**). See Supplementary Table 3 for more details.

**Extended Data Figure 9.**
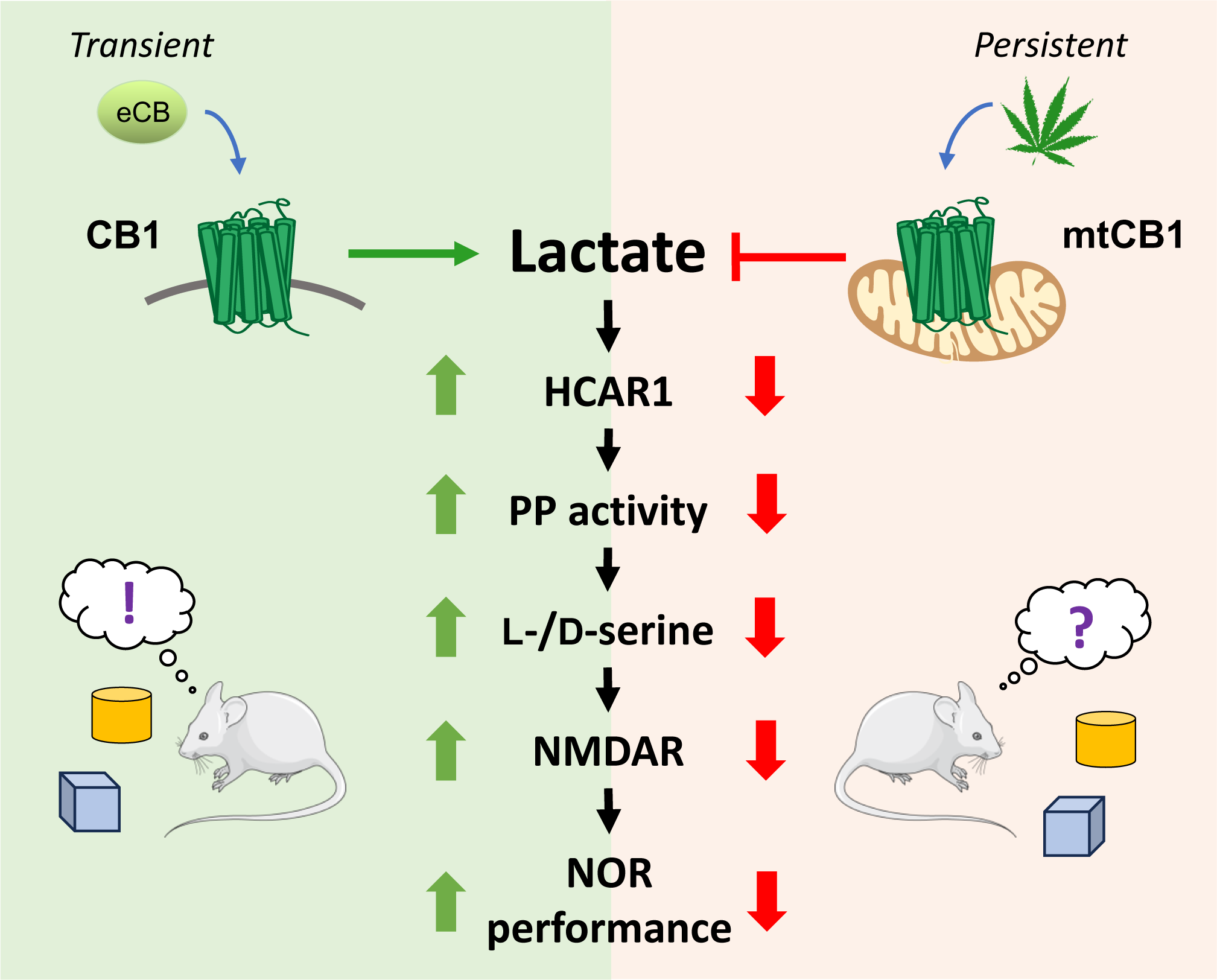
A lactate-dependent shift of glycolysis mediates synaptic and cognitive processes. Lactate promotes cognitive performance via a cascade involving HCAR1 and phosphorylated pathway (PP) activity, thereby increasing L-/D-serine levels and NMDAR activity to allow an adequate NOR memory consolidation. Importantly, whereas transient activation of non-mitochondrial CB1 receptors promote this novel lactate signaling to promote cognitive performance, the persistent activation of mitochondrial CB1 receptors impairs the lactate signaling and disrupt the consolidation of NOR memory via a specular mechanism.

## MATERIAL AND METHODS

### Animals

All experiments were conducted in strict compliance with the European Union recommendations (2010/63/EU) and were approved by the French Ministry of Agriculture and Fisheries (authorization number 3306369) and the local ethical committee (authorization APAFIS#18111). Animals used in the study were divided into two categories. The first group consisted of female and male CB1-KO, CB1-DN22 or CB1-WT mice, used as breeders to obtain newborn mice for primary cultures. The second category correspond to animals used for behavior or electrophysiology studies, consisting of male C57BL/6N (JANVIER, France), male CB1^f/f^, male GFAP-CB1-KO mutant and GFAP-CB1-WT littermate mice (two to three months-old). Animals were housed in groups under standard conditions, with free access to food and water, in a day/night cycle of 12/12 hr (light on at 7 am). Specific deletion of CB1 on GFAP positive cells in adult mice was obtained via loxP/Cre system, with a tamoxifen inducible CreERT2 recombinase^105^ encoded under the GFAP promoter^30^. Female mice carrying the ‘‘floxed’’ CB1 gene (CB1^f/f^)^106^ were crossed with CB1^f/f;GFAP-CreERT2^, to obtain CB1^f/f;GFAP-CreERT2^ and CB1^f/f^ littermates, named throughout the text GFAP-CB1-KO and GFAP-CB1-WT, respectively. For induction of CB1 deletion, 7-9 weeks-old mice were treated daily with 1 mg tamoxifen via intraperitoneally (i.p.) injections (10 mg/mL dissolved in 90% sesame oil, 10% ethanol) for 8 days. After each injection, mice were surveilled and weighted every two days to control their wellbeing. Mice were used 3-5 weeks after the last tamoxifen injection^14, 30^.

### Surgery and viral stereotaxic injection

Male CB1^f/f^ (CB1-flox) mice were anesthetized in a box containing 5% Isoflurane (Virbac, France) before being placed in a stereotaxic frame (Model 900, Kopf Instruments, CA, USA) in which 1.5% to 2.0% of Isoflurane was continuously supplied via an anesthetic mask during the whole duration of the experiment. For viral intra-hippocampal AAV delivery, mice were submitted to stereotaxic surgery and AAV vectors were injected with the help of a microsyringe (0.25 mL Hamilton syringe with a 30-gauge beveled needle) attached to a pump (UMP3-1, World Precision Instruments, FL, USA). Where specified, CB1-flox mice were injected directly into the hippocampus (HPC) (0.5 µL per injection site at a rate of 0.5 µL per min), with the following coordinates: HPC, AP -1.8; ML ± 1; DV -2.0 and -1.5. Following virus delivery, the syringe was left in place for 1 minute before being slowly withdrawn from the brain. To induce the deletion of hippocampal astroglial CB1 receptors and the rescue (RS) of CB1 receptor expression either with WT or mutant DN22 sequences, mice were injected in the hippocampus with the following combination of viral particles: (i) AAV-CAG-DIO-empty + AAV-GFAP-GFP (Control mice), (ii) AAV-GFAP-CRE-mCherry + AAV-CAG-DIO-empty (HPC-GFAP-CB1-KO mice), (iii) AAV-GFAP-CRE-mCherry + AAV-CAG-DIO-CB1-GFP (HPC-GFAP-CB1-RS mice) and (iv) AAV-GFAP-CRE-mCherry + AAV-CAG-DIO-DN22-CB1-GFP (HPC-GFAP-DN22-CB1-RS mice). Animals were used around 4-5 weeks after local AAV infusions. Mice were weighed daily and individuals that failed to regain the pre-surgery body weight were excluded from the behavioral experiments.

### Mixed cortical brain cell cultures

Mixed cortical cultures of neuronal and glial cells were prepared from 1 to 3-day-old neonatal mice as previously described^107^. Briefly, mice were euthanized, the brain removed, and cortex dissected in iced cold Hank’s balanced salt solution. The tissue was enzymatically digested with trypsin/EDTA for 5 min at 37 °C and the enzymatic digestion was stopped with 10% FBS in B-27 supplemented neurobasal medium. After this, a gently dissociation of the tissue was performed by repeatedly passing it through a 1-mL micropipette tip. Obtained cells were left in suspension to allow debris precipitation and removal. Cells were seeded in 18-mm glass coverslips treated with poly-L-lysine and incubated for 90 min to allow cell adhesion. After this, medium was replaced with fresh B-27 supplemented neurobasal medium with 10 mM glucose, 0.24 mM pyruvate, 2 mM GlutaMAX^TM,^ 100 U/mL penicillin, 100 µg/mL streptomycin and 2.5 µg/mL amphotericin B at 37 °C in a humidified atmosphere of 5% CO_2_. At day in vitro (DIV) 13–14, cultures were exposed to 1 × 10^6^ plaque forming units (pfu) of adenoviral vectors (serotype 5) coding for Laconic^17^. Measurements were carried out 48-72 h after infection of cells (DIV 16-17). Adenoviral vectors encoding the FRET biosensor were custom made by Vector Biolabs (PA, USA).

### Cell lines

HEK293T cells (ATCC, CRL-3216TM, lot 62729596) were cultured in Dulbecco’s modified Eagle’s medium (DMEM) with 1 g/L glucose, supplemented with 10% fetal bovine serum (FBS), 100 U/mL penicillin, and 100 µg/mL streptomycin, 0.1 mM of Gibco® MEM Non-Essential Amino Acids, and maintained at 37 °C in a humidified atmosphere of 5% CO_2_. For “sniffers cells” experiments, cells were transfected 18 – 24 hours before experiments with 1 µg plasmid DNA encoding for the extracellular lactate fluorescent biosensor eLACCO2.1 (Ref. 26), using polyethylenimine (PEI) as transfection agent. On the day of experiment, the DMEM media was replace for B-27 supplemented neurobasal (used for the mixed glia-neuron culture) and cells were detached gently with a micropipette. Immediately after, the cell suspension was seeded on mixed glia-neuron and incubated for 4 – 5 hours before imaging.

### Drug preparation and administration

For in vitro experiments, water soluble drugs were dissolved directly in the imaging solution (see below for its composition). Concentrated stocks of drugs prepared in DMSO were diluted directly in the imaging solution. The imaging solution contained the same amount of solvent respectively to the drug(s). For behavioral experiments, sodium lactate, 3,5-Dihydroxybenzoic (3,5-DHBA) and L-serine were prepared in saline (0.9% NaCl). Δ9-tetrahydrocannabinol (THC) and NCT-503 was prepared in a mixture of saline with 2.5% cremophor and 2.5% DMSO to obtain a solution of 0.6 mg/ml. All drugs were injected i.p. with a 26G needle, either immediately after the acquisition phase of the NOR task or after 1-hour post-acquisition phase. For injection of two drugs, a 5 min pause between injections was performed. Vehicles contained the same amounts of solvents respectively to the drug. All drugs were prepared fresh before the experiments.

### Histology

#### Mice perfusion

Mice were deeply anesthetized with an intraperitoneal injection of pentobarbital (400 mg/kg). Mice were transcardially perfused with 20 mL of phosphate-buffered solution (PBS, 0.1 M, pH 7.4) for 2 min and then with 50 mL of neutral buffered formalin 10% wt/vol (Sigma, HT501128-4L;) for 5 min. After perfusion, the brains were isolated and postfixed in neutral buffered formalin 10% wt/vol for 24 h. After this, the brains were transferred to PBS-sucrose 30% wt/vol solution for cryopreservation. Once the brains were completely dehydrated and sunk into the bottom of the tube (on average 3 to 5 days), brains were frozen in isopentane and cut in coronal sections of 30 μm using a cryostat (Leica Biosystems, CM1950S). Hippocampal slices were stored in antifreeze solution at -20°C until further use.

#### Double immunofluorescence

Free floating sections were permeabilized in a blocking solution (PBS + 10% donkey serum + 0.3% Triton X-100) for 1 hour at room temperature (RT). Then, sections were incubated overnight at 4°C with a mix of primary antibodies: chicken anti-GFAP (1:500, Abcam ab4774) and rabbit anti-GFP (1:1000, Invitrogen A11122). After several washes with PBS, slices were incubated for 2 hours at RT with a mix of secondary antibodies: goat anti-chicken Alexa Fluor 647 (1:500, Invitrogen A32933) and goat anti-rabbit Alexa Fluor 488 (1:500, Invitrogen A11008). Then, sections were incubated with 4’,6-diamidino-2-phenylindole (DAPI,1:20000, Invitrogen D3571) diluted in PBS to visualize cell nuclei (data not shown). Finally, sections were washed in PBS and mounted. Immunofluorescence images were taken with a Microscope Leica DM 4000 BLED equipped with a Camera Leica DFC 365.

### Fluorescence imaging

Mixed cortical glia-neuron cultures, HEK293T cells or a co-culture of mixed glia-neuron culture with HEK293 cells (sniffers), were mounted in an open chamber and imaged on wide-field mode with an inverted Leica DMI 6000 microscope (Leica Microsystems, Wetzlar, Germany) equipped with a resolutive HQ2 camera (Photometrics, Tucson, USA). The illumination system used was a lumencor spectra 7 (Lumencor, Beaverton, USA). The objectives used were a HC PL APO CS 20X dry 0.7 NA and a HCX PL APO CS 40X oil 1.25 NA. Multi-positions were done with a motorized stage Scan IM (Märzhäuser, Wetzlar, Germany). A 37°C atmosphere was created with an incubator box and an air heating system (Life Imaging Services, Basel, Switzerland). The system was controlled by MetaMorph software (Molecular Devices, Sunnyvale, USA). Cells were superfused with an imaging solution consisting of (in mM): 10 HEPES, 112 NaCl, 24 NaHCO_3_, 3 KCl, 1.25 MgCl_2_, 1.25 CaCl_2_, 2 glucose, 0.5 sodium lactate and bubbled with air/5% CO_2_ at 37 °C, at a constant flow of 3 mL/min. Astrocytes expressing Laconic were imaged at 40X, and excited at 430 nm for 0.01–0.05 s, emissions collected at 465-485 nm for mTFP and 542-556 nm for Venus, with image acquisition every 10 s. The ratio between mTFP and Venus was computed and normalized to the baseline. To quantify the basal lactate level (Extended Data Fig. 1A-B and Extended Data Fig. 2A-B), the biosensor occupancy was computed as a proxy of intracellular lactate level with the following equation: Occupancy = (R_0_-R_min_)/(R_max_-R_min_), in which R_0_: basal mTFP/Venus ratio (before any drug treatment), R_min_: steady state mTFP/Venus ratio induced by sodium oxamate (6 mM) or pyruvate (10 mM), R_max_: steady state mTFP/Venus ratio obtained after MCTs block (1 μM AR-C155858) or 10 mM lactate. Lactate production rates (Fig. 1 E-F) were computed by fitting a linear rate to the first minutes of lactate accumulation during MCTs block with 0.5 mM diclofenac. HEK293 cells expressing eLACCO2.1 (either alone or in co-culture) were imaged with a 20X objective, excited at 475 nm for 0.05–0.1 s and emission collected at 509-547 nm for GFP, with image acquisition every 10 s. The obtained GFP fluorescence was normalized to the baseline.

### Novel Object Recognition Memory Task

The novel object recognition (NOR) test took place in a L-shaped maze as previously described^14, 108^. The behavior task was carried out in a room adjacent to mice housing with a light intensity of 50 ± 3 lux. An overhung video camera over the maze was used to record mice behavior and scoring was performed offline. The task consisted of 3 sequential daily trials of 9 min each. On day 1, the habituation phase, mice were placed in the center of the maze and allowed to freely explore the arms in the absence of any objects. On day 2, the acquisition phase, mice were placed in the center of the maze with the presence of two identical objects positioned at the extremities of each arm and left to freely explore the maze and the objects. On day 3, the long-term memory test phase (24 hours after acquisition session), similarly to day 2, mice were placed in the center of the maze with two objects but one of the familiar objects was replaced by a novel object of different shape, color, and texture, and mice were left to explore both objects. The position of the novel object and the associations of novel and familiar were randomized. All objects were previously tested to avoid biased preference. The apparatus as well as objects were cleaned before experimental use and between each animal testing with water and at the end of experimental session with ethanol 70%. Some animals were used in two consecutive experiments: (i) First NOR task, with an injection of vehicle immediately after acquisition phase, and (ii) one week after the first NOR task, a second NOR task (with different objects) was performed, and an injection of drugs (NCT-503, THC, 3,5-DHBA, etc.) was done immediately after the acquisition phase. Cognitive performance was assessed by the discrimination index (DI), computed as the difference between the time spent exploring the novel (TN) and the familiar object (TF) divided by the total exploration time (TN+TF): DI = [TN-TF]/[TN+TF]. Object exploration was defined as the nose-poking of the objects. Mice with a total exploration time <15 s were not included in the data analysis. Memory performance was also evaluated by directly comparing the exploration time of novel and familiar objects, respectively. Experienced investigators evaluating the exploration were blind to the treatment and/or genotype of the animals. Normally, mice carried out the task without issues, however some mice performed behaviors incompatible with the test (i.e., not exploring both objects, either by chance or due peeing in a maze arm and refusing to cross over the urine.) These mice were returned to the home cage and retested one hour after. If mice failed again to perform the behavioral task, they were excluded from the experiment.

### Electrophysiology

#### Slice preparation

After being anaesthetized with 5% isoflurane for two minutes, C57BL/6N mice (12-16 weeks old) were decapitated, and the brain quickly extracted in ice-cold artificial cerebrospinal fluid (aCSF) containing (in mM): 125 NaCl, 2.5 KCl, 1 NaH2PO4, 1.2 MgCl2, 0.6 CaCl2, 26 NaHCO3 and 11 Glucose (pH 7.3, 305 mosmol/kg). Coronal hippocampal slices (350 μm) were prepared using a vibratome (Leica VT1200 S) and hemisected. Next, Slices were incubated in aCSF containing 2 mM MgCl2 and 1 mM CaCl2 during 30 minutes at 33°C. Finally, slices were allowed to rest for 1h at room temperature before starting recordings.

#### NMDAR field excitatory postsynaptic potentials recordings

Slices were transferred into a recording chamber, maintained at 32°C, and perfused continuously with aCSF (3 mL/min) containing this time 1.3 mM MgCl2 and 2.5 mM CaCl2. Extracellular field excitatory postsynaptic potentials (fEPSPs) were recorded with a Multiclamp 700B amplifier (Axon Instruments, Inc.) using pipettes (2-4 MΩ) filled with aCSF and placed in the stratum radiatum of the hippocampal CA1 area. The stimulation of the Schaffer collaterals (0.05 Hz, 100 μs duration) with a concentric bipolar tungsten electrode was used to induce synaptic responses. Recorded signals were filtered at 2 kHz and digitized at 10 kHz via a DigiData 1440 (Axon Instruments, Inc.). Data were collected and analyzed offline using pClamp 10.7 software (Axon Instruments Inc.). For each experiment, a stable baseline was recorded for at least 15 minutes before starting the Input/Output measurement. The stimulation amplitude that evokes 50% of the maximal response was used for recordings. NMDA-fEPSPs were then isolated with low Mg^2+^ aCSF (0.2 mM) in the presence of 2,3-dihydroxy-6-nitro-7-sulfamoyl-benzo[f]quinoxaline-2,3-dione (NBQX, 10 μM) to block AMPA/kainate receptors, respectively. To study the co-agonist binding site occupancy of synaptic NMDARs, after obtaining a 20-minute stable response, D-serine (50 μM) was bath applied for 30 minutes. The same protocol was applied with lactate (2 mM). For occlusion experiments, the first drug was bath applied 20 min before the recording and then during the whole experiment. Drug used were D-serine (50 μM), lactate (2 mM), the PHGDH inhibitor NCT-503 (10-20 μM) and the HCAR1 agonist 3,5-DHBA (1 mM). D-AP5 (50 µM) was applied at the end of 3,5-DHBA occlusion experiments to confirm that NMDA-fEPSPs were mediated by NMDA receptors (Extended data Fig. 7A,B). NMDAR-fEPSPs slopes were measured as a linear fit of the rising phase, between time points set in the baseline period and corresponding to 20% and 60% of the peak amplitude. The change in slope was normalized to the baseline slope taken during the 15 min immediately before drug applications. Representative NMDAR-fEPSPs traces are the average of successive sweeps that correspond to the last 10 minutes of baseline versus the last 10 minutes of drug applications.

### Quantification and statistical analysis

All graphs, linear regressions and statistical analyses were performed using GraphPad software (version 8 or 10). Data is presented as time course (average of imaged cells ± SEM, from a representative experiment), scatter plots of individual cells and bars + symbols. In some experimental data, the SEM is small enough to be contained inside symbols. Otherwise, time courses without error bars correspond to a representative cell from an independent experiment and is indicated in the relevant figure legend. All data was analyzed for outliers using the ROUT method (false discovery rate of 1%), and for normality with the Shapiro Wilk test. To perform the statistical analysis, the electrophysiology data was averaged every 5 min. Differences between groups were assessed using the average of each independent experiment using either by paired t-test, unpaired t-test, one-way ANOVA followed by Tukey’s multiple comparison test, Kruskal-Wallis followed by Dunn’s multiple comparison test, Mixed-effects model followed by Dunnett’s multiple comparison test or two-way ANOVA followed by Tukey’s multiple comparison test. P< 0.05 was considered significant and are annotated in each figure and available on Supplementary Table 2 and 3. Otherwise, nonsignificant data is indicated as NS.

---

**Suppl. Table 2.**
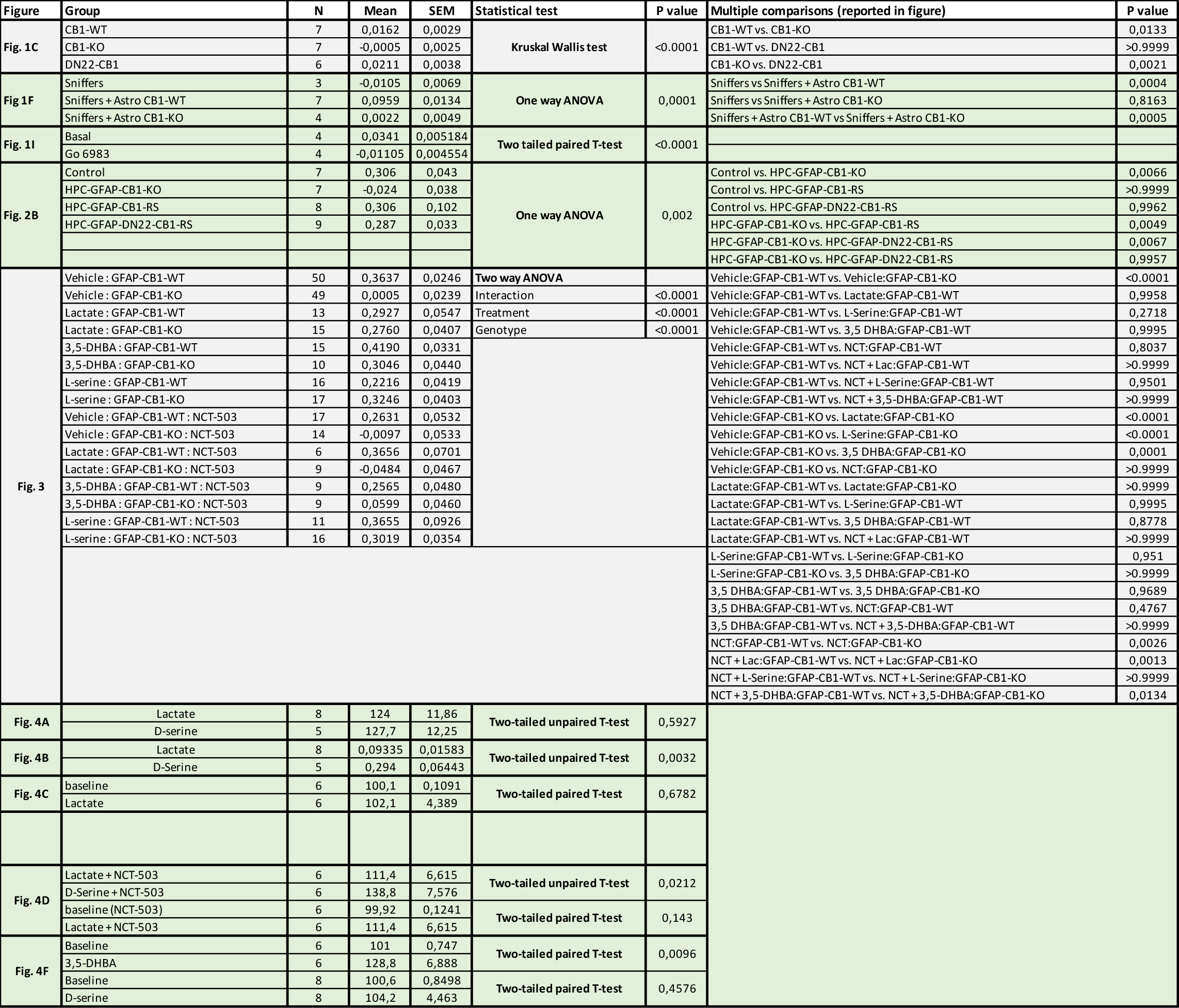
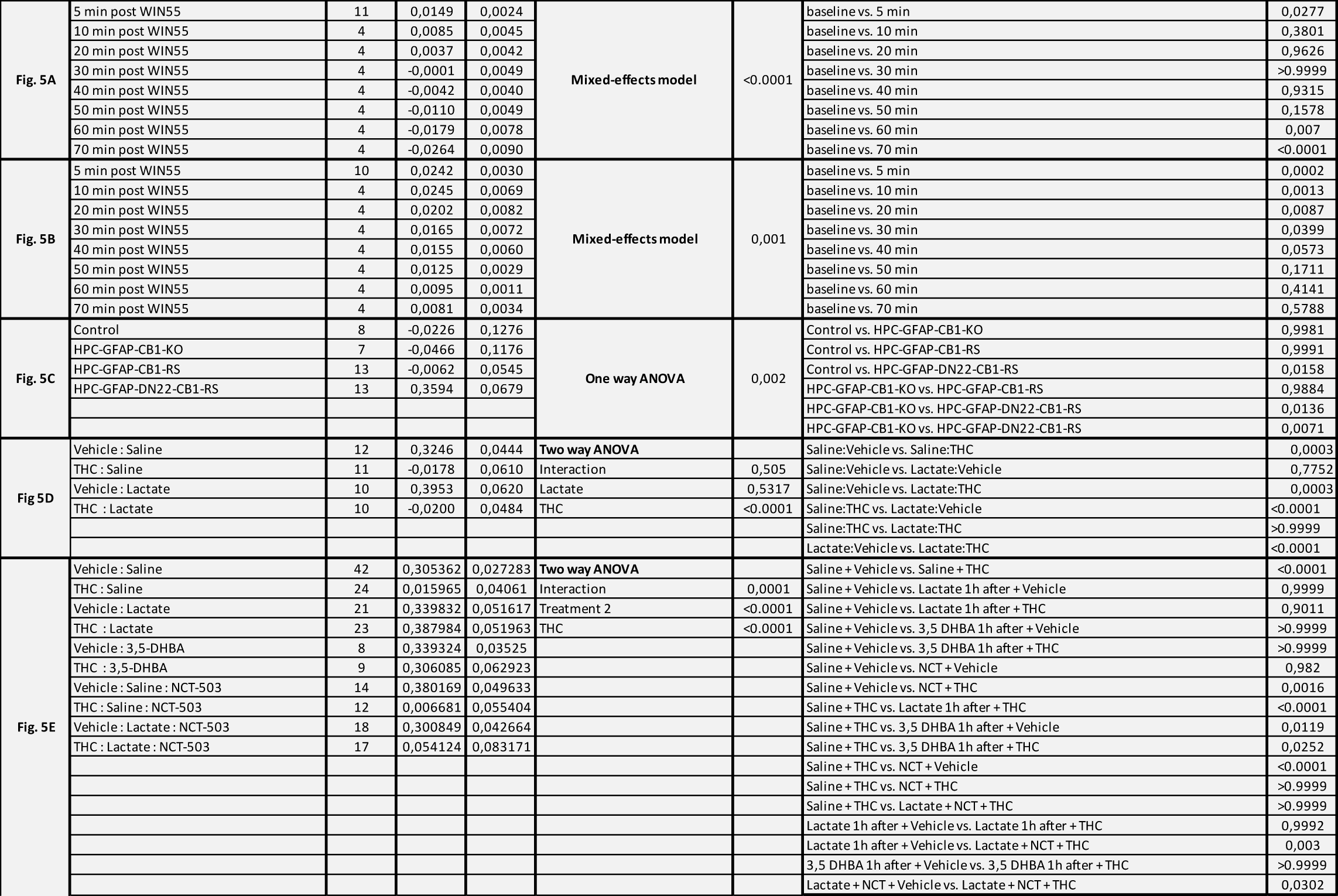

**Suppl. Table 3.**
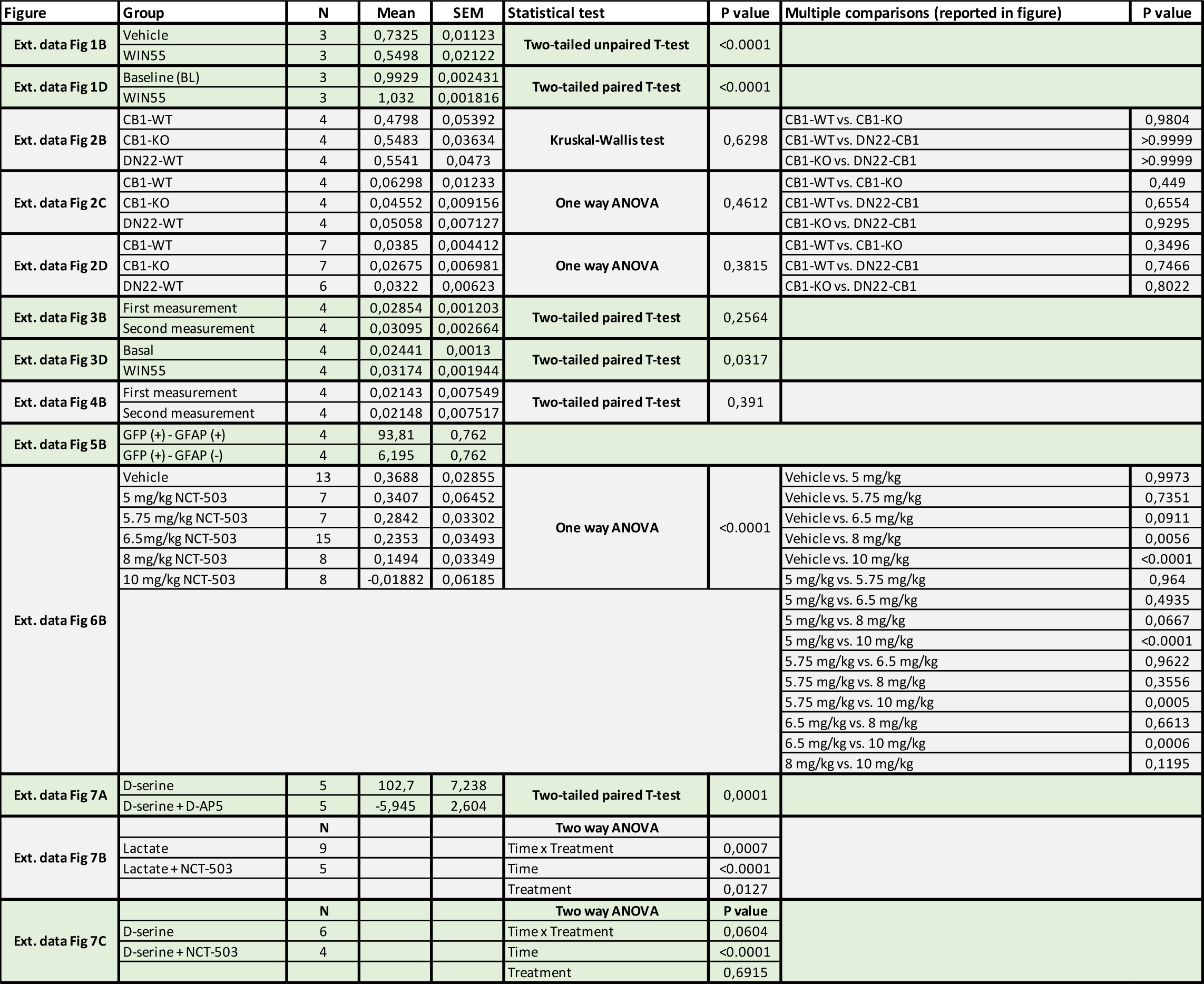

